# G34R cancer mutation alters the conformational ensemble and dynamics of the histone H3.3 tails

**DOI:** 10.1101/2025.05.02.651932

**Authors:** Harrison A. Fuchs, Yunhui Peng, Shine Ayyapan, Houfang Zhang, Ruben Rosas, Anna R. Panchenko, Catherine A. Musselman

## Abstract

In recent years, mutations in the histone variant H3.3 have been discovered in pediatric and adult gliomas and osteosarcomas. One of these is the G34R mutation in the H3.3 N-terminal tail. While this mutation is known to disrupt epigenomic pathways, the effects on nucleosome structure itself have not been explored. In light of recent studies, that have demonstrated that the interaction of the H3 tail with nucleosomal and linker DNA is driven in large part by arginine residues, we sought to determine if the G34R cancer mutation and adjacent G33R mutation, not observed in cancer, directly alter nucleosome structural dynamics. Using NMR spectroscopy and molecular dynamics simulations, we investigate the effects of these mutations on the H3 tail in the context of the nucleosome. Our results show that both of these mutations enhance the association of the H3 tails with DNA and decrease the conformational dynamics around the site of mutation. Moreover, our results reveal that both mutations lead to changes in the conformational ensemble of the entire tail, re-positioning it on the nucleosomal DNA as well as promoting intra-tail interactions. This has implications for the accessibility of the H3 tail to effector proteins and for changes in higher-order chromatin structure resulting from a shift in intra-versus inter-nucleosome contacts mediated by the H3 tail.

## INTRODUCTION

The eukaryotic genome is compacted into the cell nucleus in the form of chromatin, a complex of the DNA with histone proteins. The core histone proteins H2A, H2B, H3, and H4 form an octameric core around which ∼147 base pairs of the DNA wrap to form the basic subunit of chromatin, the nucleosome core particle (NCP). Each of the histones has a canonical form as well as several variants. While the canonical histones are expressed in a replication dependent manner, variants are expressed throughout the cell cycle and act as replacement histones, that are placed in response to a specific event or at a specific genomic region. In addition to the canonical versions of H3 (H3.1 and H3.2), there are several H3 variants including H3.3. As a replication-independent variant histone, H3.3 is one of the most highly conserved histones and found to be enriched at dynamic genomic regions such as gene promoters, active gene bodies, and cis regulatory elements. However, it is also found at telomeres and heterochromatic regions (1). The H3.3 variant differs from H3.1 and H3.2 at only 5 and 4 amino acids respectively (Supplementary Figure 1a): one of these is located in the N-terminal tail, where the canonical A31 is substituted by serine, while the other amino acids are located in the core at the H3.3 specific chaperone interface. Notably, phosphorylation at the H3.3 specific S31 residue has an important functional role at enhancer regions (2).

The H3.3 protein is known to be encoded by the cancer driver gene (3). Moreover, it was reported that ∼30% of pediatric gliomas contained mutations in H3.3, including those at residue G34 (H3.3G34V/R) in the H3 tail (4). Further studies have also found the G34R mutation in adult gliomas and osteosarcomas (4–8). Work in yeast has revealed recombination defects and genomic instability in the presence of H3.3G34R, and a mouse model revealed impaired DNA damage repair and activation of the cGAS/STING pathway (9). Fission yeast, expressing only the H3G34R mutant, demonstrated a global reduction in tri-methylation and acetylation at H3K36 (10). H3.3G34R leads to a loss of H3K36me2/3 in cis (i.e localized to the mutated histone) and inhibited SETD2 activity in vitro (11–13). The loss of H3K36 methylation has further been found to impair recruitment of the DNA methyltransferase DNMT3A (12). In contrast, another study reported preferential binding to and inhibition of the KDM4 demethylase at H3.3G34R leading to an increase in H3K9 and H3K36 methylation across the genome (14, 15). The H3.3G34R mutation was also found to inhibit binding of the H3K36me3 reader ZMYND11 (16), whereas ZMYND8 was seen to preferentially bind the mutated H3.3G34R tail (17). Thus, H3.3G34R can dramatically alter the epigenome through direct alteration of enzyme (writer/eraser) or reader domain activity.

Here we explore additional mechanisms, namely if the H3.3G34R mutation can alter chromatin through directly modulating the nucleosome conformational dynamics and solvent accessibility of histone tails. Previous work from our laboratories and others have demonstrated that the H3 tails interact with DNA in the nucleosome context (18–20). Globally the tail is thought to be in a primarily DNA-bound state adopting a broad ensemble of conformations spanning various DNA binding configurations from the two nucleosome gyres to the dyad. On the other hand, locally tail residues are thought to transition quickly between multiple non-distinct DNA sites. Together, this is consistent with a concept of a so-called fuzzy complex (21, 22).

Tail-DNA interactions have been shown to be driven in large part by arginines and lysines (18, 19, 23). The H3 tail residues can generally be grouped into two segments enriched in lysine/arginines which are flanked by flexible TGG motifs (Figure 1a). The TGG containing element closest to the NCP core (comprising T32, G33, and G34) is the most flexible region of the H3 tail and has been proposed to act as a hinge allowing for switching between dyad and two gyre-bound conformational ensembles(20, 23–25). We thus set out to test if mutations of these hinge glycines to arginines might alter the conformational ensemble and/or dynamics of the H3 tail. Using a combination of molecular dynamics (MD) simulations and nuclear magnetic resonance (NMR) spectroscopy, we examine the oncogenic G34R mutation along with the adjacent G33R and find alterations in both the conformational ensemble and dynamics of the tail. These changes are observed throughout the hinge region, but intriguingly propagate through the H3 tail.

**Figure 1.**
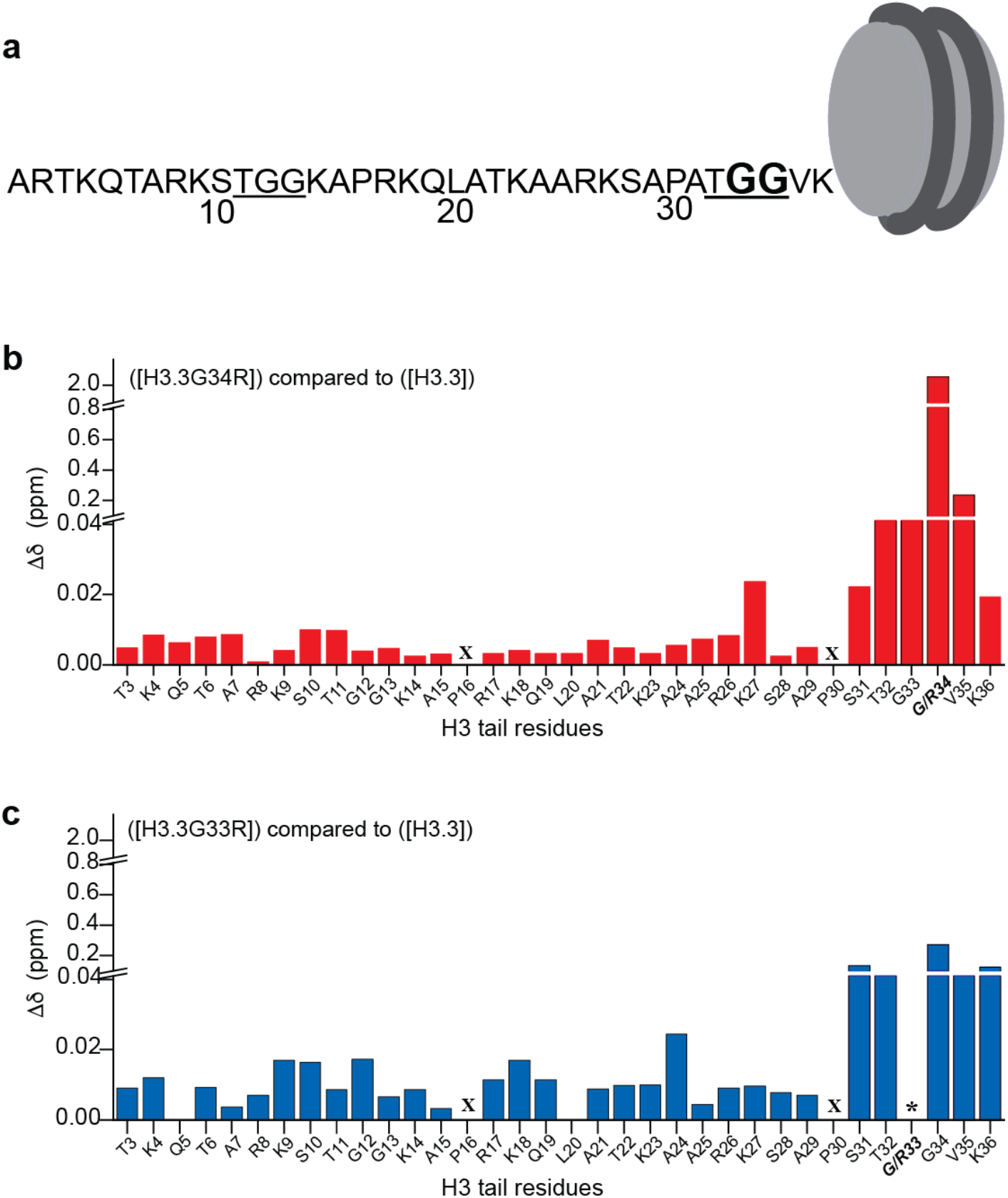
Hinge mutations alter the conformational ensemble of the entire H3 tail. **a)** The H3.3 tail sequence with both TGG elements underlined. Glycine 33 and 34 mutated in this study are in bold. **b, c)** Normalized chemical shift changes (Δδ) in ^1^H,^15^N-HSQC spectra are plotted as a function of H3.3 tail residues between WT and H3.3G34R containing nucleosomes (b, red) or WT and H3.3G33R containing nucleosomes (c, blue). X denotes a proline residue which does not have a backbone amide and thus no corresponding peak, * denotes a missing peak. All data were collected at 100mM NaCl.

## METHODS

### Histone protein preparation

The histone H3.3 coding sequence (Uniprot accession P84243) with the common C110A and G102A mutations was obtained from Genscript (Piscataway, NJ) and placed in a pET3a vector. The Q5 mutagenesis kit (New England BioLabs) was used to generate plasmids with either the G33R or G34R mutation. Human histones H2A (Uniprot accession P0C0S8), H2B (uniprot accession P62807), and H4 (Uniprot accession P62805) were also encoded in pET3a vectors. H2A, H2B, and H4 were overexpressed in either Rosetta 2 (DE3) pLysS or BL21 (DE3) E. coli in 2xYT media. Histone H3.3 and mutants were overexpressed in BL21 (DE3) E. coli in M9 minimal media enriched with vitamin (Centrum), 1 g/L ^15^NH_4_Cl, and 5 g/L D-glucose or 3 g/L of ^13^C-D-glucose. Cells were grown to an OD_600_∼0.4, induced with IPTG (0.2 mM for H4 and 0.4 mM for H2A, H2B, or H3), and incubated at 37°C for 3 hours for H3 and H4, or 4 hours for H2A and H2B. Histones are then purified out of inclusion bodies as described previously (26). Briefly, bacterial cells are lysed via sonication, and after removing the soluble portion of the lysate, the histones were resuspended from the cell pellet in 6 M Guanidine-HCl. The samples are then exchanged into 8 M urea for the purpose of further purification through ion exchange chromatography, first using Q Sepharose Fast Flow followed by a step gradient over SP Sepharose Fast Flow resin (Cytiva Life Sciences). Purified histone proteins are then exchanged out of urea into water to avoid unwanted carbamylation. After exchange into water, Electrospray ionization mass spectrometry was used to validate the histones and ensure no carbamylation occurred during purification. These defined proteoforms with no modifications will be referred to with the [] notation, e.g. [H3.3] (27).

### Preparation of Widom 601 DNA

The 147 bp Widom 601 DNA was amplified in DH5a E. coli (New England Biolabs) using a plasmid containing 32 repeats of the Widom 601 sequence for 16 hours at 37°C. Cells were harvested, and nucleic acid was extracted using alkaline lysis methods, followed by precipitation using isopropanol. DNA was resuspended in 10 mM Tris-HCl pH 8.0, 50 mM EDTA and incubated with RNAse overnight at 37°C to degrade all cellular RNA. Protein was then purified out of the sample using phenol extraction, followed by removal of phenol with 24:1 chloroform:isoamyl alcohol. The plasmid DNA was then precipitated away from the small RNA fragments by polyethylene glycol (PEG) precipitation by addition of NaCl and 6K PEG to final concentrations of 0.5 M and 10% w/v respectively. The precipitated DNA was then resuspended in 10 mM Tris-HCl pH 8.0, 0.1 mM EDTA buffer and subjected to another round of phenol extraction followed by a chloroform:isoamyl alcohol cleanup. The 147 bp Widom 601 sequence was then excised from the purified plasmid DNA through incubation at 37°C with EcoRV-HF restriction enzyme (New England Biolabs) over the course of several days (until complete as determined by agarose gel). The plasmid was purified away from the 601 DNA by PEG precipitation adding PEG 6K to a final concentration of 7.4% w/v and NaCl to a concentration of 0.525 M. The DNA was then purified by ion exchange chromatography using a Resource Q column (Cytiva Life Sciences). DNA concentration as calculated using UV-Vis Spectroscopy and the extinction coefficient of 2,312,300.9 M^-1^ cm-1.

### Histone octamer formation

The refolding of histone proteins into octamer was carried out as detailed previously (28). Briefly, equimolar amounts of the unfolded purified histone proteins were resuspended in 20 mM Tris-HCl pH 7.5, 6 M Guandidne-HCl, 10 mM DTT, mixed, and then refolded by dialysis into 20 mM Tris-HCl pH 7.5, 2 M KCl, 1 mM DTT, 1 mM EDTA, 0.5 mM benzamidine hydrochloride at 4°C. After refolding, octamers were purified by size exclusion chromatography using a HiLoad 16/600 Superdex 200 pg column (Cytiva Life Sciences). Purified octamers were analyzed by 18% SDS-PAGE gel and concentrations determined by UV-Vis spectroscopy.

### Reconstitution of nucleosome core particles

Purified histone octamers were mixed with the purified 147 bp Widom 601 DNA in a molar ratio of 1:1.2 in 20 mM Tris-HCl pH 7.5, 2 M KCl, 1 mM DTT, 1 mM EDTA, 0.5 mM benzamidine hydrochloride in a volume that brought the total concentration of DNA to 0.7 mg/mL. The mixture was desalted via dialysis to a final concentration of 150 mM KCl using a linear gradient over approximately 48 hours at 4°C. Samples were dialyzed against 20 mM Tris-HCl pH 7.5, 1 mM DTT, 1 mM EDTA, 0.5 mM benzamidine hydrochloride. Concentrated nucleosomes were heat shocked for 30 min at 37°C to ensure optimal positioning. Samples were then purified by running them over a 10-40% sucrose gradient using an SW-32 rotor and spinning them at 32K rpm for 45 hrs. Nucleosome formation was confirmed through sucrose gradient profile, 5% Native PAGE gel and 18% SDS PAGE gel (Supplementary Figure 1b). Nucleosome concentrations were determined by UV-Vis spectroscopy using diluted nucleosome samples dissociated in 2M salt and the extinction coefficient for the 601 DNA. Reconstituted nucleosomes will be referred to with a ([]) notation, e.g. ([H3.3]) indicates a nucleosome containing the H3.3 variant, canonical H2A, H2B, and H4, with all residues unmodified (27).

### NMR spectroscopy

All experiments were carried out on a Bruker AVANCE NEO 600 MHz spectrometer equipped with a cryoprobe.

*Assignments:* Resonances for ([H3.3]), ([H3.3G33R]), and ([H3.3G34R]) were largely transferred from the ([H3.2]) assignments performed previously (BMRB ID 50806). These were confirmed by collection of an HNCACB on 110-170 μM ([^13^C,^15^N-H3.3]), ([^13^C,^15^N-H3.3G33R)], or ([^13^C,^15^N-H3.3G34R]) samples at 45°C in 20 mM MOPS pH 7.0, 1 mM EDTA, 1 mM DTT, and 10% D2O. Data was processed with NMRpipe (29) and analyzed with CcpNMR Analysis (30).

*Chemical shift analysis:* ^1^H,^15^N-HSQC were collected on ([^15^N-H3.3]) (105-130 mM), ([^15^N-H3.3G33R]) (140-187 mM), and ([^15^N-H3.3G34R]) (180-235 μM) at 37°C in a sample buffer containing 20 mM MOPS pH 7.0, 1 mM ETA, 1 mM DTT, 10% D2O, and 0mM or 100mM NaCl. Chemical shift perturbations between constructs were calculated as:

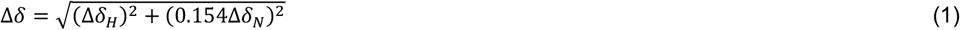

where Δδ_H_ and Δδ_N_ are the differences in the ^1^H and ^15^N chemical shift, respectively, for each resonance between samples.

^*15*^*N-relaxation analysis* Longitudinal (R_1_) and transverse (R_2_) ^15^N relaxation rates were measured on ([^15^N-H3.3]) (105-130 μM), ([^15^N-H3.3G33R]) (140-187 μM), and ([^15^N-H3.3G34R]) (180-235 μM) at 37°C in a sample buffer containing 20 mM MOPS pH 7.0, 1 mM ETA, 1 mM DTT, 10% D2O, and either 0mM or 100mM NaCl. TopSpin pulse sequences hsqct1etf3gpsi3d and hsqct2etf3gpsitc3d were utilized with 2,048 (^1^H) and 200 (^15^N) total points, acquisition times of 106 ms (^1^H) and 51 ms (^15^N), and spectral widths of 16 ppm (^1^H) and 32 ppm (^15^N).

For 0mM NaCl, R_1_ experiments were collected with relaxation loop lengths of 10 ms, 50 ms (x2), 100 ms, 200 ms, 400 ms, 500 ms (x2) and 800 ms, 1,000 ms, 1,250 ms (x2), and 1,500 ms. The R_2_ experiments used total relaxation CPMG loop lengths of 15.68 ms, 31.36 ms (x2), 47.04 ms, 78.4 ms, 94.08 ms (x2), 109.76 ms, 125.44 ms, 156.8 ms, 188.16 ms (x2), and 203.4 ms. For 100mM NaCl, R_1_ experiments were collected with relaxation loop lengths of 25 ms, 200 ms, 400 ms (x2), 800 ms, 1,250 ms, and 1,750 ms. The R_2_ experiments used total relaxation CPMG loop lengths of 15.68 ms, 31.36 ms, 47.04 ms (x2), 62.72 ms, 94.08 ms (x2), 109.76 ms.

Spectra from nuclear spin relaxation experiments (^15^N-R_1_, and ^15^N-R_2_) were processed with NMRpipe (29). The R_1_ and R_2_ relaxation rates were determined by least-square fitting of the amide peak intensities to a single-exponential decay without offset. The R_2_/R_1_ values were calculated by dividing the respective fit values. The rotational correlation time, *τ*_*c*_, was calculated from R_1_ and R_2_ using the following equation

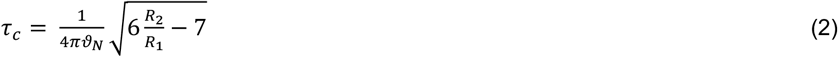

Where P_*N*_ is the resonance frequency of ^15^N. The errors in R_1_ and R_2_ were propagated to R_2_/R_1_, and then to *τc* using standard error propagation.

### Thermal Stability Shift Assay

Reconstituted NCPs were diluted to a final concentration of 2.25 µM in thermal stability assay buffer [20 mM Tris-HCl pH 7.5, 0.5 mM TCEP and 0 mM, 100 mM, or 150 mM NaCl]. The SYPRO orange dye (Sigma-Aldrich) was added to each sample at a final concentration of 5X in a 20 µL reaction volume. The fluorescence from the dye was monitored in a CFX Opus 384 real-time PCR system (Bio-Rad) from 25 ºC to 95 ºC with incremental steps of 1 ºC. After each temperature was reached, the samples were incubated for 1 minute prior to measurement of the fluorescence signal. At least three replicates for each nucleosome sample and negative control (only buffer) condition were prepared and measured. The raw fluorescence values of the SYPRO dye obtained from the real-time PCR machine (RFU) were normalized as:

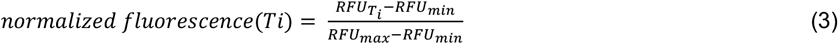

where RFU_Ti_ is the raw fluorescence at the temperature i, RFU_min_ and RFU_max_ are the lowest and highest raw fluorescent values, respectively, obtained for each sample among the entire range of temperatures measured.

To determine the melting temperature (Tm) associated with nucleosome denaturation transition steps, we plotted the rate of change of the normalized fluorescence signal (d(normalized fluorescence)/dT) as a function of temperature. The rate of change was calculated as:

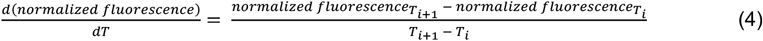

For plotting d(normalized fluorescence)/dT vs temperature, the resulting d(normalized fluorescence)/dT value was assigned to the Ti+1 temperature. In this distribution, the temperature at each peak indicates the nucleosome melting temperature at nucleosome melting transition steps.

### Construction of nucleosome models containing wild-type and mutant H3 tails

The human homotypic nucleosome structural models were constructed based on the high-resolution X-ray crystal structure (PDB ID: 1KX5), where histones were substituted by human variant H3.3 and human canonical H2A, H2B and H4 with Widom 601 DNA sequence. The simulated system was exactly the same as used in our experiments. G33R and G34R mutations were introduced into two copies of H3.3 tails time using Chimera (31). As a result, three models were obtained: H3.3, H3.3G33R, and H3.3G34R nucleosomes. Each model was used as an initial conformation for molecular dynamics (MD) simulations. In an initial conformation the histone tails were linearly extended into the solvent symmetrically oriented with respect to the nucleosomal dyad axis. To prevent the DNA fraying at the terminus of DNA molecules, we applied harmonic constrains to the terminal nucleotides to limit the maximum distance between O4-N6 and N3-N1 atoms to less than 150% of the initial distance in the PDB structure.

### Molecular Dynamics Simulations

All MD simulations were performed with the Amber 22 package (32) using the AMBER ff14SB force field for protein and OL15 for double-stranded DNA (33, 34) and OPC water model (35). The nucleosome models were solvated in a cubic water box with at least 20 Å from the nucleosome atoms to the edge of the water box. The NaCl from the12-6 HFE parameter set for monovalent ions were added up to 150mM. Systems were maintained at T = 310 K using the Langevin thermostat with collision frequency γ = 2 ps^-1^. The Berendsen barostat was used for constant pressure simulation at 1 atm. SHAKE bond length constraints were applied for bonds involving hydrogens and Hydrogen Mass Repartitioning was applied to allow a 4 fs integration time step to be employed (36). The cut-off distance for non-bonded interaction calculations was 8 Å. Particle Mesh Ewald (PME) method with a grid point spacing of 1 Å and real space cut-off of 12 Å was applied for the electrostatic calculations. Periodic boundary conditions were used, and the trajectories were saved every 20 ps. The solvated systems were first subjected to 10,000 steepest descent minimizations and then for another 10,000 conjugate gradient minimizations. Then, the systems were gradually heated from 100 to 310 K in the NVT ensemble and then subjected to 2 ns equilibration before production runs. Each simulation was performed for 2 µs for each system in NPT ensemble with five independent simulation runs.

### Analysis of simulation trajectories

Trajectories of each simulation run were visualized and analyzed using a set of customized TCL and Python scripts that utilized the capabilities of VMD (37), 3DNA (38), and AMBER Tools (32). The trajectory frames were firstly superimposed onto the initial structure by minimizing RMSD values of Cα atoms in histone cores. The first 200 ns frames of each simulation trajectories were excluded from the analysis. The tail–DNA atomic interactions were defined between two non-hydrogen atoms from DNA and histone within a distance of less than 4 Å, and atomic contacts were calculated for trajectory frames of every 1 ns. In-house codes were developed to quantify the DNA and histone paths, tail-DNA contacts and radius of gyration. The binding free energy between histone tails and DNA was calculated using the molecular mechanics generalized Born surface area (MM/GBSA) method implemented in the Amber 22 package (32). We performed calculations for every 1 ns frame (ignoring the first 50 ns), and residue-wise decomposition was applied to derive the binding energy per tail residue.

### Calculation of NMR observables from the MD trajectories

The NMR spectroscopy parameters of histone tails were calculated from the MD trajectories using our recently developed package based on the spectral density approach (39–41) (manuscript in preparation). Briefly, the autocorrelation function *C(t)*, which describes relaxation due to dipole-dipole interactions between the ^1^5N and ^1^H atoms of protein backbone amide groups, was directly calculated from the coordinate trajectory obtained from molecular dynamics (MD) simulations. Then, the spectral density, *J(ω)*, was obtained by transforming the correlation function into the frequency domain via Fast Fourier Transform (FFT).

In our calculation of autocorrelation function, all MD frames were aligned to reference coordinates (initial structure) via Cα atoms of the globular histone fold to eliminate the overall tumbling motion of nucleosome. Then, the effects of overall motion were reintroduced by multiplying the calculated correlation function by *exp(−τ/τ*_*rot*_), where *τ*_*rot*_ is rotational correlation time of the NCP assumed to be 163.4 ns (42). To improve the convergence of the results, we have split the 2000 ns long-time MD trajectory into 20 chunks with equal length and calculated the averaged *C(t)* values over all chunks. The autocorrelation functions were fitted with three-exponentials to derive the spectral density and NMR spectroscopy parameters. This approach allows us to calculate the NMR observables, including relaxation rates R_1_ and R_2_, and the Nuclear Overhauser Effect (NOE), considering factors like Larmor frequencies, gyromagnetic ratios, NH bond length, and chemical shift anisotropy. Next, the residue specific rotational correlation time, *τ*_*c*_, was calculated using equation (2)(43). The calculated NMR parameters were quantitively compared with our experimental data, including R_1_, R_2_ and *τ*_*c*_ values for H3 tail residues.

### Calculation of residue-residue interaction networks

We analyzed histone H3 residue-residue interaction network using RING 4.0 (44) which generated residue-residue interaction networks from our MD trajectories and categorized interactions into hydrogen bonds, disulfide bridges, ionic, Van der Waals, π–cation and π–π interactions. Interactions are represented as edges with an associated probability score (the fraction of MD trajectories in which that interaction is present). The structural ensemble of WT, G33R and G34R H3 tails from simulations were subjected to the RING 4.0 webserver for analysis with default parameters. Trajectory frames were extracted every 10 ns from each run (five runs of 2000 ns long), and the conformations of two copies of H3 tails were combined, resulting in a total of 2000 conformational states for each ensemble.

## RESULTS

### NMR CSPs reveal that mutation of the hinge alters the H3 tail conformational ensemble

To test if the cancer mutation G34R alters the conformational ensemble of the H3 tail, we used NMR spectroscopy. Nucleosome core particles were reconstituted with [^15^N-H3.3] wild type (WT) or mutant [^15^N-H3.3G34R], and unlabeled [H2A], [H2B], and [H4] using the 147 bp Widom 601 sequence. Here the brackets denote a defined histone proteoform, in this case confirmed to contain no modifications. Reconstituted nucleosomes are denoted with parenthesis notation, e.g. ([^15^N-H3.3]) which denotes a nucleosome containing [^15^N-H3.3] and where all other histones are assumed to be canonical (27) (see methods).

Heteronuclear single quantum coherence (HSQC) spectra were recorded on ([^15^N-H3.3]) or ([^15^N-G34R]) in buffer containing either 0 mM NaCl or 100 mM NaCl. Analysis of the ^1^H,^15^N-HSQC spectra (referred to as WT and G34R spectra hereafter in the text) revealed that a single resonance was observed for each tail residue despite there being two tails, supporting that the two H3 tails in each NCP are structurally identical or nearly identical within the nucleosome context (Supplementary Figure 2,3). In the G34R spectrum a strong peak was observed for R34 near the other arginine resonances (Supplemental Figure 3a). Comparison of spectra for WT and G34R revealed several chemical shift perturbations (CSPs). The largest CSPs are observed in and around the putative hinge region itself, including residues T32, G33, V35, and K36 (Figure 1b). This is due at least in part to the chemical environment change arising from the mere presence of an arginine side chain rather than a glycine side chain, but also reports on resulting changes in the structural ensemble of the tail. Interestingly, in addition to these CSPs we also observed smaller CSPs in distal residues along the entire length of the histone tail. Together, these perturbations indicate a change in the conformational ensemble of the entire H3 tail upon mutation of G34 to arginine. Similar observations were made in either 0 mM or 100 mM NaCl, though the signal in 100 mM NaCl was substantially weaker overall (Figure 1, Supplementary Figures 2-4).

To investigate if these changes are specific to the mutation of G34R or are generic to mutations of the TGG hinge (i.e. either G33R or G34R) we also generated and tested ([^15^N-H3.3G33R]). Similar to the G34R spectrum, we saw a single set of peaks for all H3 residues in the G33R spectrum. Notably, we were unable to identify a resonance for R33, likely due to conformational exchange on the intermediate timescale (Supplementary Figure 2,3). Also, similar to G34R, CSP differences between the G33R and WT spectra were the largest in and around the site of mutation, but were also observed along the entire length of the H3 tail (Figure 1c). Notably, the magnitude of CSPs in distal residues were observed to be larger due to the G33R mutation as compared to the G34R mutation.

Taken together, chemical shift analysis suggests that mutation of either G33 or G34 to arginine alters the conformational ensemble of the entire H3 tail but in a manner unique to the position of mutation. In other words, the two glycines in the TGG hinge have distinct effects on structural properties of the H3 tail.

### NMR relaxation analysis reveals that mutations of the hinge reduce the H3 tail conformational dynamics

To assess if G33R and G34R mutations alter the conformational dynamics of the tail, we measured ^15^N spin relaxation rates R_1_ and R_2_ for ([^15^N-H3.3]), ([^15^N-H3.3G33R]), and ([^15^N-H3.3G34R]) (see Supplementary Figures 5-8). Data were again collected at 0 mM and 100 mM NaCl, with 0 mM concentration leading to substantially better signal and more complete data sets. From these relaxation rates the residue-specific effective correlation times (τ_C_) were calculated. Comparisons with WT reveals that mutation of either glycine to arginine leads to an increase in the τ_C_ values of residues in the hinge indicating a decrease in conformational flexibility of this region (Figure 2). In contrast to observations via CSPs there was no consistent change in the τ_C_ values for residues distal to the mutations. Although an increase in τ_C_ values of K14-L21 was observed for G34R at 0mM NaCl (Supplemental Figure 7), this was not recapitulated at 100mM NaCl, nor was it observed for G33R.

**Figure 2.**
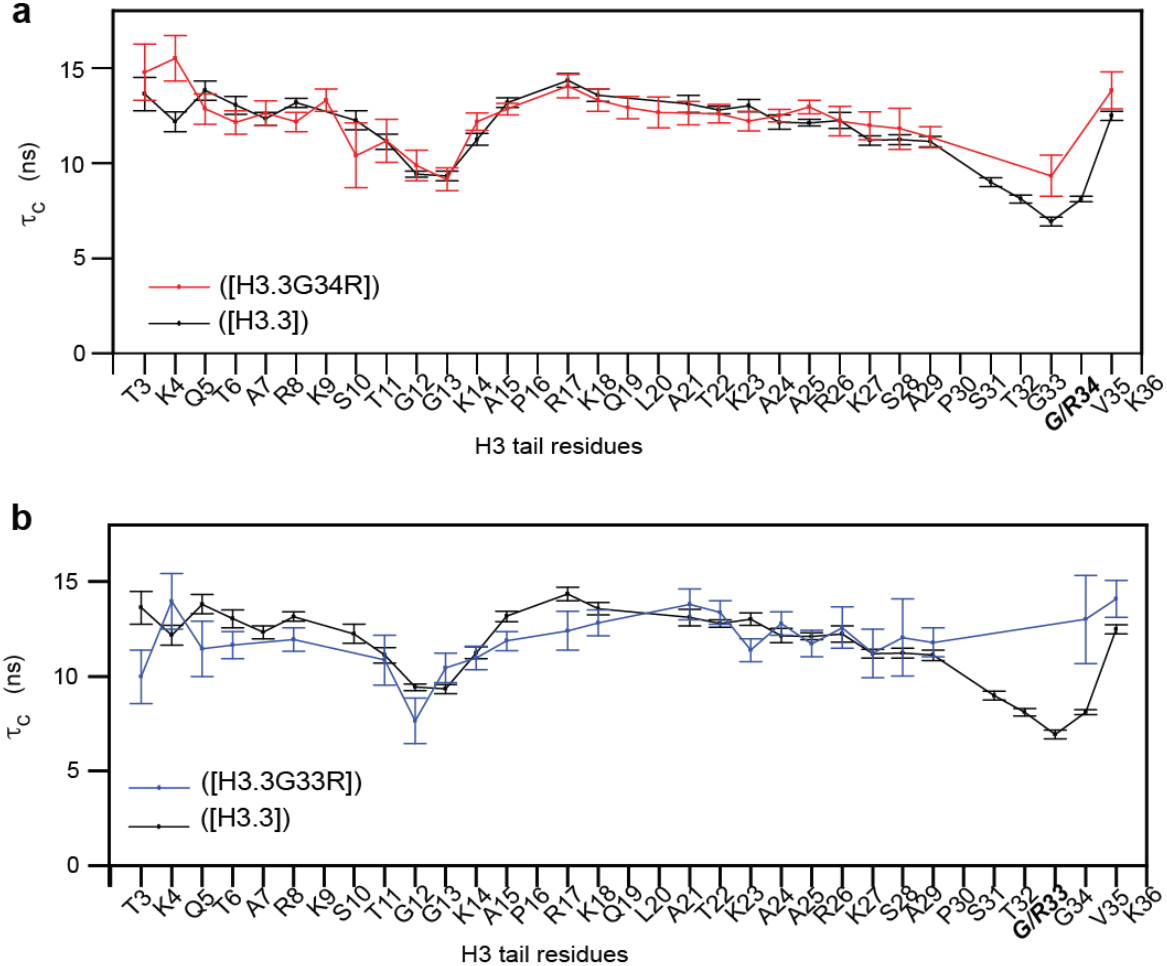
Hinge mutations reduce localized dynamics. Amino acid specific overall correlation times (τ_C_) as a function of H3.3 tail residue for WT nucleosomes (black) or mutants H3.3G34R containing nucleosomes **(a**, red) or H3.3G33R containing nucleosomes **(b**. blue). Error bars were calculated as described in the Methods section.

Altogether, this indicates that G33R and G34R stabilize the TGG hinge in the H3 tail, whereas little to no change in the dynamics of the distal tail residues were observed at least at near physiological salt concentrations. This change in dynamics is accompanied by changes in the conformational ensemble of the entire tail as denoted by changes in the CSPs (see previous section).

### Global stability of the NCP does not change upon mutations of the hinge

The H3 tail has previously been shown to contribute to the thermal stability of the nucleosome (45). To determine if these mutations have an impact on the overall stability of the NCP, we analyzed WT and mutants using a thermal stability shift assay (46). At 0 mM NaCl, WT and both mutants showed a biphasic nucleosome disassembly profile with melting temperatures of ∼72 ºC (first peak) and ∼81 ºC (second peak) (Figure 3a). The appearance of two unfolding transition steps has been interpreted as the uncoupled dissociation of the H2A/H2B dimers (first peak) and the H3/H4 tetramer (second peak) and is observed for some histone variant nucleosomes subjected to this assay (46, 47). The G33R mutation demonstrated a very moderate shift to higher temperatures relative to WT and G34R with both melting temperatures increasing by approximately ∼1 ºC suggesting slightly increased stability (Figure 3a). However, at 150mM or 100mM NaCl, we observed no differences between WT and mutants. At these near physiological salt concentrations, a single major transition step was observed with a substantially attenuated (as compared to that observed for 0mM NaCl) second step. No differences were observed in the disassembly profile or major step melting temperature between WT and mutants, which were measured to be 70 ºC at 150 mM and 71 ºC at 100 mM NaCl (Figure 3b, c). (Note that the attenuated second step was too small to be able to measure an associated melting temperature.) Together, these results indicate that while the G33R and G34R mutations decrease the conformational dynamics of the H3 tails and alter the conformational ensemble, they do not affect the overall stability of the nucleosome core particle under near physiological conditions.

**Figure 3.**
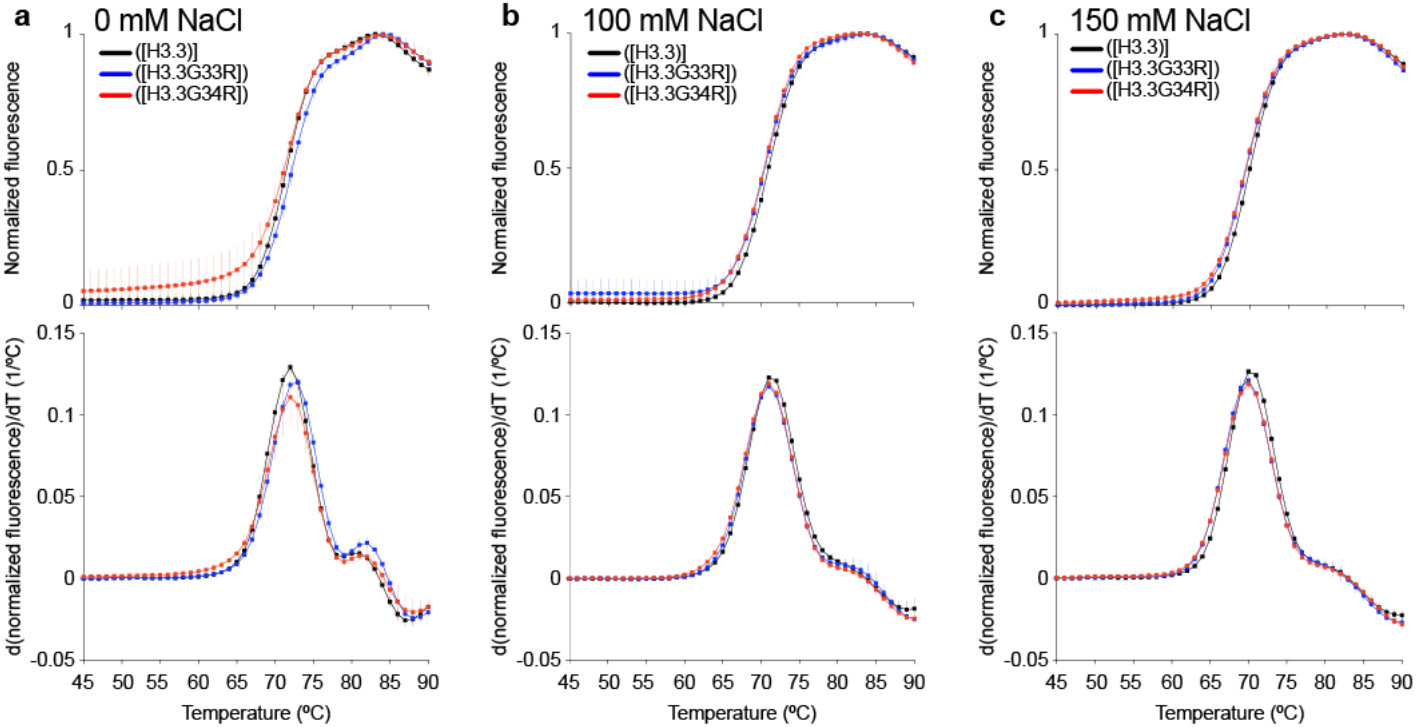
Hinge mutations do not substantially alter the overall stability of the H3.3 NCP. Thermal stability shift assays of WT (black), G33R containing (blue), and G34R containing (red) nucleosomes in 0mM NaCl **(a)**, 100mM NaCl **(b)**, or 150mM NaCl **(c)**. Top plots show the thermal denaturation profile of nucleosome samples by plotting the normalized SYPRO orange signal as a function of temperature. Bottom plots show the differential values of the thermal stability curves presented at the top to reveal the inflection points in the thermal stability curve, which correspond to the two NCP transition unfolding steps. Data from at least three experiments were plotted as the mean and standard deviation (error bars).

### Computational analyses show decreased tail dynamics and increased tail-DNA interactions for hinge mutants

To assess the atomistic basis of the observed changes we turned to MD simulations. In total, five all-atom MD simulation runs (2μs each) were carried out for each of H3.3, H3.3G33R, and H3.3G34R nucleosome systems. Using the raw trajectories from the simulations of H3.3, H3.3G33R, and H3.3G34R nucleosomes, we have calculated the R_1_, R_2_ and *τ*_*c*_ values per tail residue and quantitively compared our predictions with experimental values (Figure 4 and Supplementary Figure 9, 10). In general, the R_2_ rates were overestimated by a factor of three, compared to experiments, suggesting that the tails are either over-stabilized in simulations or that the trajectories are not long enough to sample the full conformational ensemble. However, the trend of computed R_1_, R_2_ and *τ*_*c*_ values agreed well with experimental values, yielding a very high Pearson correlation coefficient for *τ*_*c*_ of up to R = 0.82 at 100 mM NaCl concentration (Figure 1 and Supplementary Table 1). Importantly, this indicates that the internal dynamics across the H3 tails is well represented in the MD simulations.

**Figure 4.**
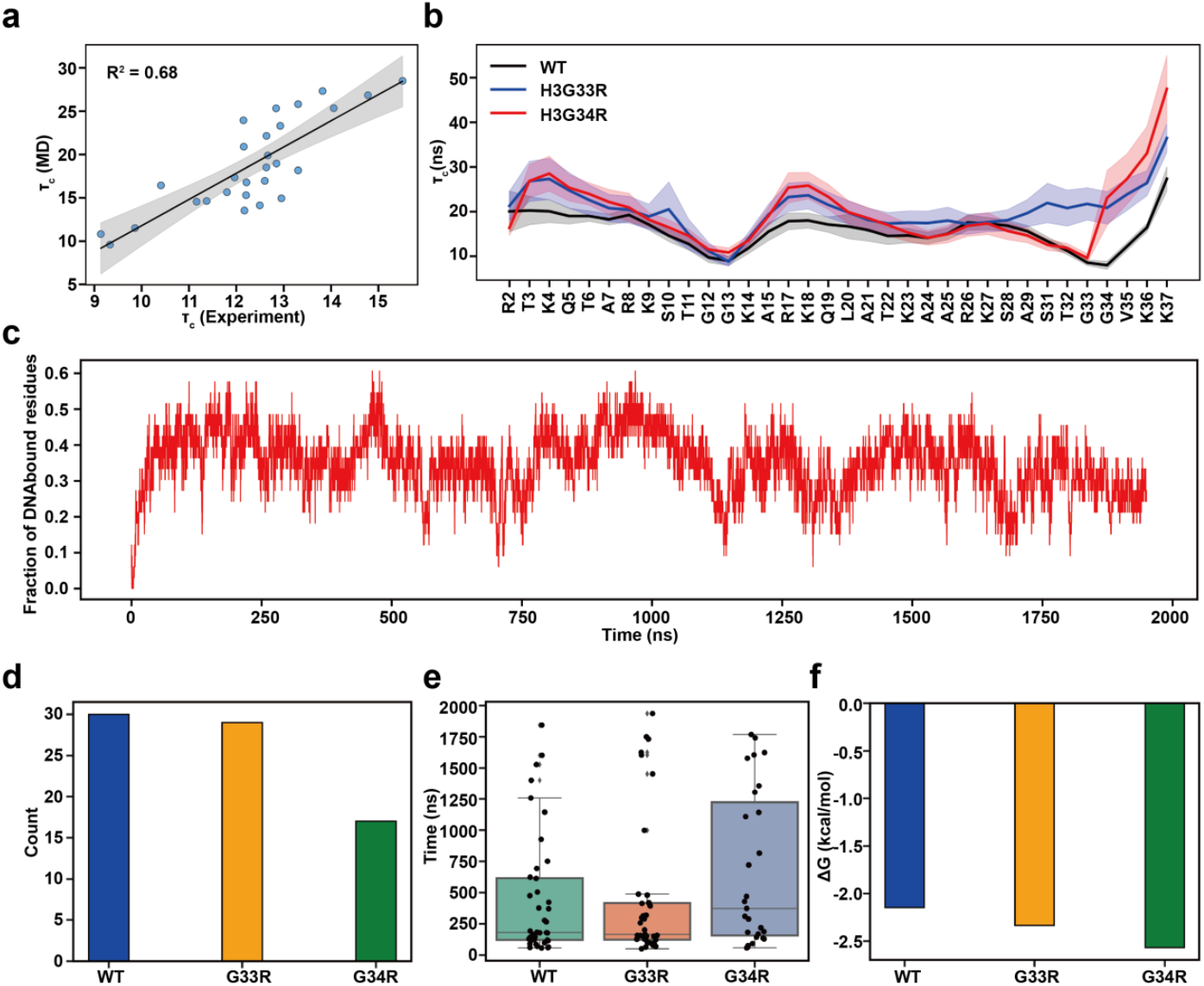
Dynamics and tail-DNA interactions of wild-type and mutant H3 tail within the nucleosome. **(a)** Comparison of the MD predicted and experimental rotational correlation times (τ_c_) for H3.3 tails at 100mM NaCl. The reported values are averaged for WT tails calculated from independent MD runs or both histone copies (n=10). b) Comparison of the rotational correlation time (τ_c_) of the motions between wild-type (WT) and mutated H3 tails. The shaded area represents the standard errors of the mean, calculated from independent simulation runs. c) A representative run shows a fraction of DNA bound residues and tail binding/unbinding events during simulation for wild-type H3 tails. d) A total number of full histone tail binding/unbinding events observed in all simulations for both copies of histones. e) Full histone tail residence time. Each point represents a binding/unbinding event observed in simulations for each histone copy. Data points with residence time shorter than 50 ns were excluded as this time is required for establishing stable interactions with DNA. An unbound state for the full tail is defined if no more than 10% of the tail residues maintain contacts with DNA. Box-plot elements are defined as: center line, median; box limits, upper and lower quartiles; whiskers are drawn at values equal to 1.5× interquartile range. f) The histone tail-DNA standard binding free energy derived from counting the number of bound/unbound events in histone tail conformational ensemble in all simulations for both copies of histones: 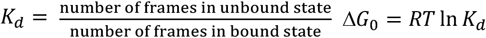

Note that since simulations were performed at the physiological NaCl concentration of 150mM, they agreed best with the experimental results obtained at 100 mM. The effect of mutations on the computed relaxation rate values also showed a very similar trend as was observed in experiment (Figure 4 and Supplementary Figure 9 and 10). Namely, both mutants led to decreased τ_C_ values compared to WT around the site of mutation.

In addition, we have also calculated the number of tail-DNA contacts and the standard binding free energies using the MM/GBSA approach. Our results indicate that both G33R and G34R mutations lead to significant increases in binding free energies of H3 tail-DNA interactions and reduced flexibility, with the G34R mutation exerting a stronger effect (Supplementary Figure 11). The largest effects on binding for the G34R mutant are seen for residues located near the mutated sites, with smaller but significant increases in tail-DNA binding also observed for residues distal to mutations, especially for residues T3-R8 R17-K18, R26-K27 and K36 (Supplementary Figure 12). Interestingly, the effect of the two hinge mutations on the R26-K27 region is opposite, with the G33R mutation decreasing the number of contacts with DNA in this region.

We further characterized the binding kinetics of H3 tail-DNA interactions and identified many rapid interconversions between tail–DNA bound and unbound states, highlighting the intrinsically dynamic nature of histone tail behavior (Figure 4b and Supplement Figure 13). To better understand the impact of mutations on H3 tail binding kinetics, we quantified the total number of transitions from unbound to bound states and calculated the residence time of the entire histone tails on DNA molecules (Figure 4 and Supplementary Figure 14). Our results show that the G34R mutation significantly increases the residence time of H3 tails on DNA molecules and reduces the number of unbinding events (Figure 4). Lastly, we estimated the dissociation constant and standard binding free energy of the tail-DNA interaction based on histone tail conformational ensemble statistics, which revealed enhanced H3 tail–DNA binding, with a more pronounced effect observed for the G34R mutant (Figure 4 and Supplementary Table 2). These results are consistent with our MM/GBSA calculations described above. We should mention that absolute values of binding energy between H3 tail and DNA estimated from histone tail conformational ensemble statistics (Supplementary Table 2) were somewhat larger in amplitude in our previous study(19), likely due to the inclusion of linker DNA regions in the simulated system, which formed a substantial number of tight contacts with the H3 tails. Additionally, the longer simulation time in the previous study allowed for a more comprehensive sampling of tail conformations with larger residence time.

### Hinge mutations enhance intra-tail interactions, promoting compact tail conformations

According to the fuzzy complex model, that is currently emerging as the dominant model of the tail dynamics in the context of the nucleosome, the tail-DNA contacts have been of primary focus. While the most recent studies on the H3 tails have indicated a lack of any secondary structure (20, 48, 49), compaction onto the DNA in principle provides the potential for intra-tail contacts. To our knowledge these intra-tail interactions have not, as yet, been thoroughly investigated. Here we analyzed the histone-histone residue networks within the H3 tail residues using the RING 4.0 server (44). The interaction graph was generated using the MD conformational ensembles of WT, G33R and G34R H3 tails (Figure 5, Supplementary Table 3). Residue-residue interaction networks were constructed by accounting for hydrogen bonds, disulfide bridges, ionic interactions, Van der Waals forces, π–cation, and π–π interactions. Each interaction was represented as an edge with an associated probability score, indicating the fraction of MD trajectories in which that interaction was observed.

**Figure 5.**
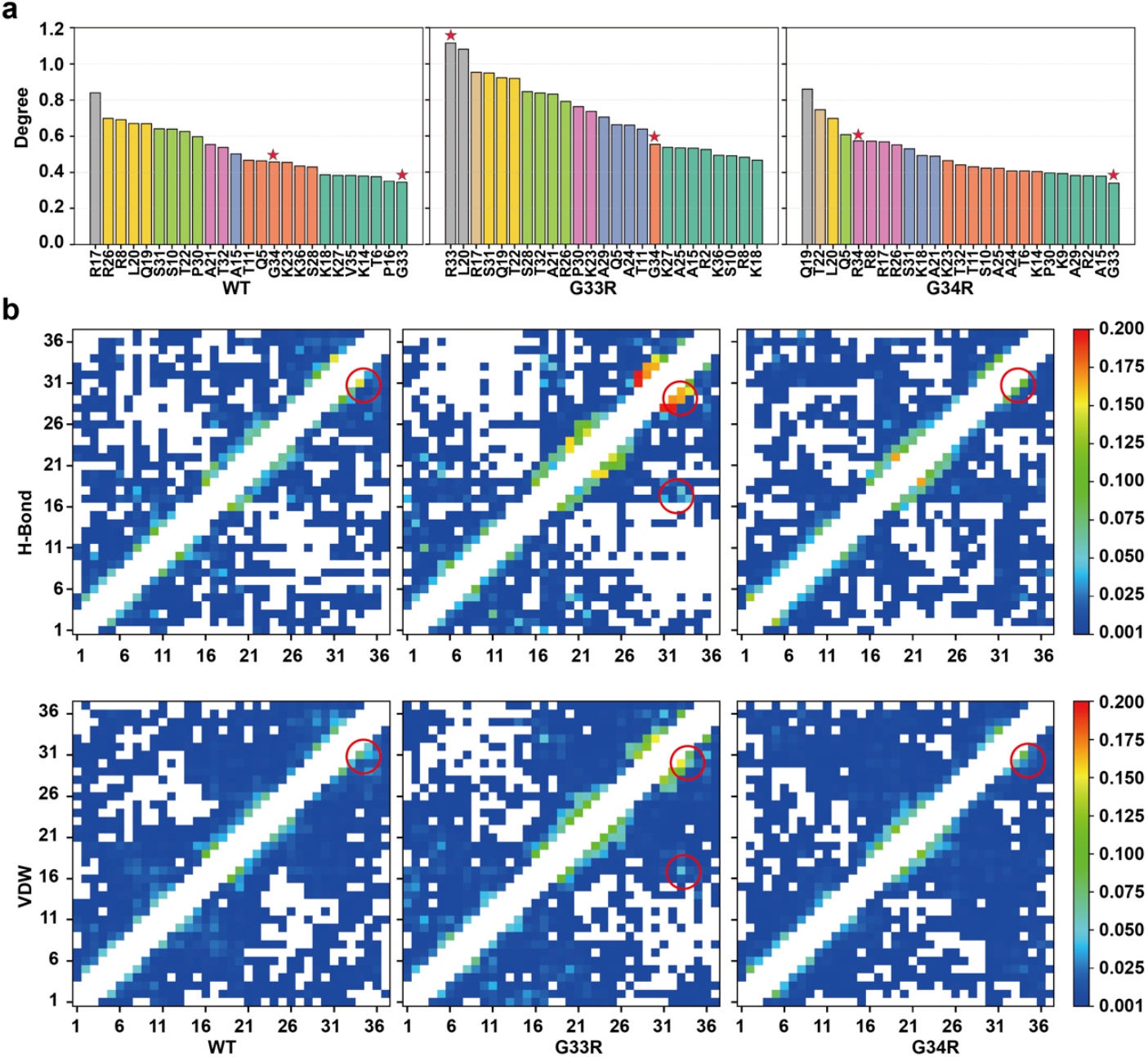
Analysis of the long-range residue communication networks from the MD simulation trajectories. a) Rank of the degree of nodes (tail residues) from the interaction graph generated from the structural ensembles in simulations using the RING 4.0 sever. The node degree is calculated by summing the probability score of all edges originating from this node. The nodes with similar node degree values are shown in same color and the G33R and G34R mutation sites are highlighted with asterisks. Nodes with high node degrees refer to tail residues with many non-covalent interactions with other residues. b) The contact map shows for each residue (x-axis) the probability of being in contact with any other residue (y-axis) via hydrogen bonds (H-bonds) and Van der Waals (VDW) interactions. The interactions between hinge region (G33 and G34) and other tails residues were highlighted with red circles. White regions indicate that no interactions were observed between the tail residues. Diagonal regions, corresponding to interactions between adjacent residues, were ignored.

Notably, the WT tail demonstrated arginine residues (R8, R17, and R26) having the highest probability scores (calculated as the sum of probability scores for all interactions involving a residue) of the intra-tail interaction network. In the G33R mutant, the cumulative probability score of R33 significantly increased compared to wild-type G33 (from 0.34 to 1.12), making it a highly connected node among all H3 tail residues. Similarly, in the G34R mutant, the node degree of R34 increased as compared to WT G34, though much less than R33, from 0.46 to 0.58. Notably, the G33R mutation led to a substantial increase in the long-range interactions between the hinge region (residues 33 and 34) and other tail residues, whereas the impact of G34R mutation was relatively minor (Figure 5b). The most significant enhanced interactions were observed between R33 and P30, R33 and A29, R34 and S31, R33 and K36, T32 and S28, as well as between G33 and P30 (Supplementary Table 4). These findings indicate that both mutations enhance intra-tail interactions, with the G33R mutation exerting a much stronger effect, highlighting the distinct roles of these two glycine residues in the hinge region. This result is supported by the differences in the radius of gyration. Both G33R and G34R mutations were found to induce more compact tail conformations compared to the WT tails, although the effect size is small (Supplementary Figure 15). Together, these results suggest that both mutations enhance intra-tail interactions in H3 histone tails, promoting more compact tail conformations.

### Hinge mutations alter the tail-DNA binding modes

We demonstrated that both G33R and G34R mutations significantly change the binding affinities and kinetics of H3 tail-DNA interactions. To further investigate whether these mutations also affect the binding modes of H3 tails on nucleosomal DNA, we analyzed the preferred binding regions by quantifying the mean number of nucleosomal DNA contacts per DNA base pair for WT and mutant H3 tails. We also evaluated changes in contact numbers induced by the mutations (Figure 6 and Supplementary Figure 16, 17). Our results reveal that both G33R and G34R mutations enhance H3 tail-DNA interactions near the dyad regions compared to the WT, with the G34R mutation exerting a stronger effect than G33R (Figure 6). Conversely, interactions of the H3 tail with the DNA regions at SHL +/-6 to +/-7 (the two DNA gyres) are significantly reduced in both mutants relative to WT. This likely explains why mutations do not substantially increase the overall stability of the NCP (Figure 3). While the tail does have increased interactions with the DNA, which would promote stability, the fact that these additional contacts are primarily at the dyad, rather than the entry/exit DNA, means that they would not have a large effect on unwrapping. In addition to H3 tail rearrangements, both mutations also led to rearrangements of H2A and H2B tails on nucleosomal DNA. For example, interactions of the H2A C-terminal tails with SHL +/-6 to +/-7 were reduced in the mutated H3 tails, while the binding of H2B to SHL +/-4 to +/-6 was significantly increased in the G33R mutant (Figure 6).

**Figure 6.**
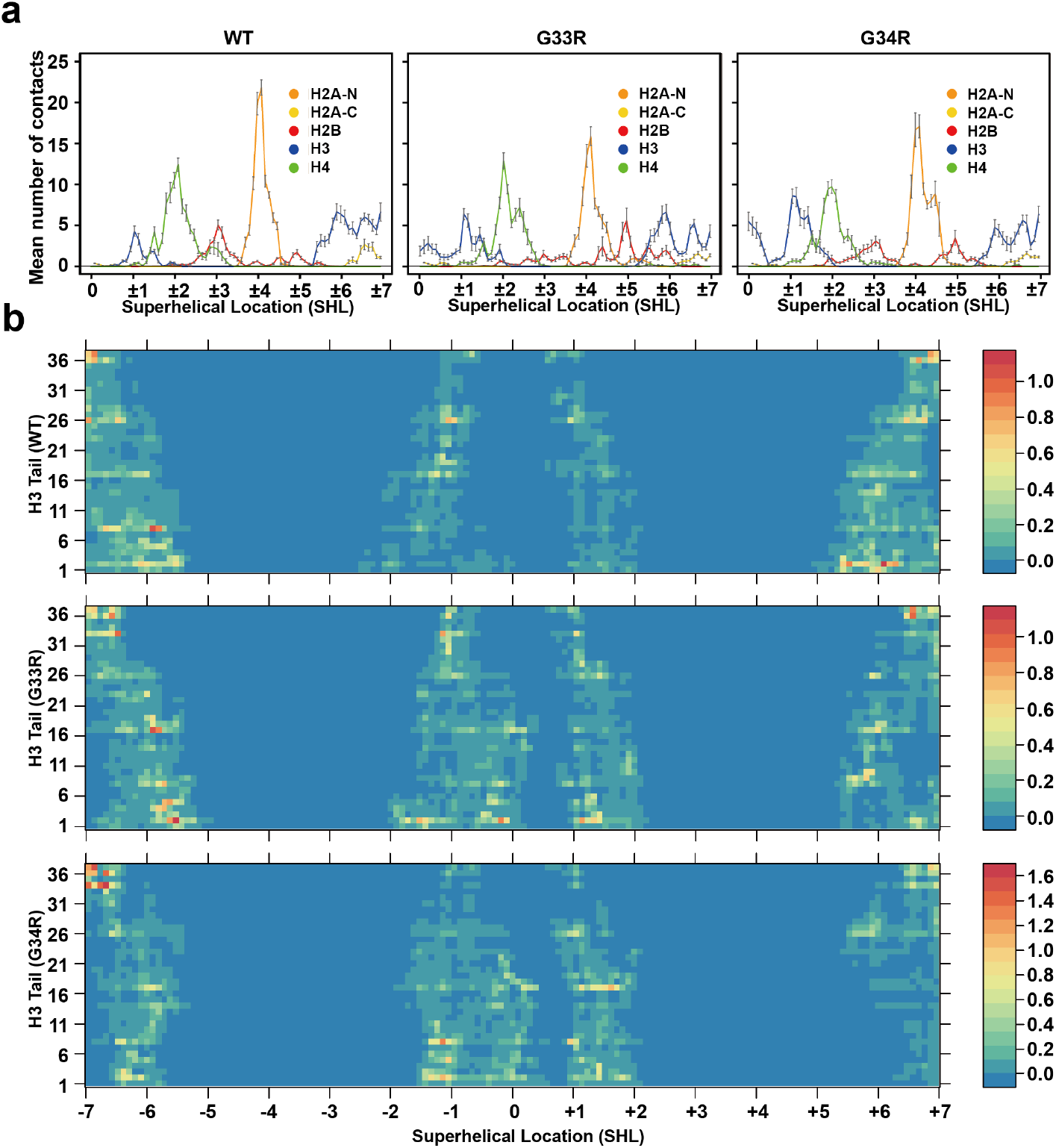
H3 G33R and G34R mutations modulate histone tail–DNA binding modes and DNA accessibility. a) Mean numbers of nucleosomal DNA contacts with WT and mutant tails per DNA base pair. The reported values are averaged and the error bars represent the standard errors of the mean for WT and mutant tails calculated from independent MD simulation runs (n=10). b) A heat map of mean numbers of contacts between each DNA base pair and tail residues for WT and mutant tails.

## DISCUSSION

A molecular model of the H3 tail dynamics in the context of the nucleosome is emerging in which the tails adopt a so-called high-affinity fuzzy conformation with the nucleosomal DNA (21, 22, 42). In this model, the tails are primarily interacting with the DNA through arginine and lysine residues distributed along the length of the tail. While the exact conformations adopted are still not fully clear, substantial evidence is mounting that the tails can associate with linker DNA, double helical gyres and the dyad. To adopt all of these conformations requires a large rearrangement of the tail on the surface of the nucleosome. Notably, there are two TGG elements in the tail that act as dynamical hotspots (see Figure 1) and the one closest to the nucleosome core has been proposed to act as a hinge for movement of the tail between the gyres and the dyad (20, 23–25). Recent studies have identified that mutation of H3.3G34 to arginine in this putative hinge region is a potent cancer driver mutation. While several studies have revealed that this mutation may directly disrupt binding to chromatin co-factors, especially those targeting H3.3K36 (50), we wondered if this could alter the conformational ensemble and dynamics of the tail itself.

In this study we utilize a combination of NMR spectroscopy and MD simulations to demonstrate that H3.3G33R or H3.3G34R mutations alter the conformational ensemble and dynamics of the H3.3 tail in the context of the nucleosome. The NMR data support that both mutations lead to a decrease in the conformational dynamics of H3.3 residues S31-K36 and alter the conformational ensemble of the entire tail. Computational analysis gives us insight into the atomistic details underlying these changes revealing: 1) an increased number of tail-DNA interactions especially around the additional arginine residue, 2) increased intra-tail interactions and a more compact tail overall, and 3) altered relative population of the tail engagement on the dyad versus double helical gyres. Altogether this supports the previous hypothesis that the region containing G33 and G34 acts as a hinge for the global dynamics of the H3 tail between the dyad and double helical gyres. However, the analysis presented here also reveals a highly complex conformational landscape of the H3 tails that include both tail-DNA and intra-tail contacts.

Previous studies have shown that the H3 tail conformation in the context of the nucleosome can inhibit the placement and readout of histone post-translational modifications (PTMs), and that tail acetylation and phosphorylation can enhance accessibility readers and writers (18, 48, 51, 52). The Morrison lab also recently demonstrated that mutations of key arginine residues along the H3 tail enhance its conformational dynamics (23). Here we observe the converse effect, where increasing the number of arginines by way of mutation decreases conformational flexibility of the tail through enhanced interactions with DNA as well as intra-tail contacts. Indeed, both mutations lead to a decrease in the conformational dynamics and tighter tail-DNA binding of H3.3, especially for residues S31-K36. This can, in turn, reduce the accessibility of these functionally important residues (like S31 and K36 which can carry PTMs). In fact, our previous studies as well as those of others have shown that the level of conformational dynamics correlates with tail accessibility and thus we expect this would inhibit the accessibility of factors targeting the hinge region.

Our results also reveal that the effect of these mutations may potentially propagate down the full length of the tail. For instance, G34R leads to enhanced tail-DNA interactions for H3R8, R17 and K18 (Supplementary Figure 3), and G33R leads to the doubled number of tail-DNA contacts for H3K9. Such long-range effects have previously been observed, including our recent study on the effects of histone tail PTMs on tail dynamics (19, 23, 48, 53). It is becoming clear that the conformational ensemble and dynamics of the tail residues are correlated, which has strong implications for regulation of the epigenome through indirect effects of nucleosome structure.

It is known that histone mutations may alter the DNA repair pathways and contribute to the carcinogenesis (54). H3.3G34V leads to altered recruitment of MutSα, a mismatch repair (MMR) protein, whereas H3.3G34R/K mutants are defective in homologous recombination and exhibit elevated rates of chromosome loss. Altogether these results support that beyond the direct effects of H3.3G34R on recruitment, this cancer mutation will have indirect effects on recognition through tail occlusion as well as direct effects on chromatin structure. The exact nature of the latter will need to be investigated in future studies but could include general decrease in tail accessibility to acting factors, altered chromatin remodeling activity, and changes to higher order structure due to a shift in intra-versus inter-nucleosome contacts by the H3 tail.

