## Supplementary file for "G34R cancer mutation alters the conformational ensemble and dynamics of the histone H3.3 tails"

| Tail Type | Experimental condition | NMR observables | Correlation Coefficient |
|---|---|---|---|
| WT | 0 NaCl, H3.3 | $R_1$ | 0.36 |
| | | $R_2$ | 0.69 |
| | | $\tau_c$ | 0.70 |
| | 100mM NaCl H3.2 | $R_1$ | -0.10 |
| | | $R_2$ | 0.77 |
| | | $\tau_c$ | 0.80 |
| G33R | 0 NaCl, H3.3 | $R_1$ | 0.12 |
| | | $R_2$ | 0.32 |
| | | $\tau_c$ | 0.30 |
| | 100mM NaCl H3.2 | $R_1$ | 0.05 |
| | | $R_2$ | 0.45 |
| | | $\tau_c$ | 0.48 |
| G34R | 0 NaCl, H3.3 | $R_1$ | 0.13 |
| | | $R_2$ | 0.51 |
| | | $\tau_c$ | 0.54 |
| | 100mM NaCl H3.2 | $R_1$ | 0.05 |
| | | $R_2$ | 0.72 |
| | | $\tau_c$ | 0.82 |

**Supplementary Table 1.** The Pearson correlation coefficients between the calculated relaxation rates from MD simulations and those measured experimentally by NMR for H3 tails in the nucleosome containing wild type H3.3 (WT), or H3.3G33R or H3.3G34R mutants.

| H3.3 tail | Number of frames in unbound state | Number of frames in bound state | Kd(M) | ∆G(kcal/mol) |
|---|---|---|---|---|
| WT | 1442 | 47313 | 0.0305 | -2.15 |
| G33R | 1291 | 57215 | 0.0226 | -2.34 |
| G34R | 742 | 48013 | 0.0155 | -2.57 |

**Supplementary Table 2.** Estimating the dissociation constants (Kd) and standard binding free energy (ΔG) for the histone tail interaction with DNA from histone tail conformational ensemble statistics. The histone tail unbound state is defined if the percentage of tail residues maintaining contacts with the DNA molecule is no more than the 10%. The dissociation constant ( $\mathbf{K_d}$ ) can be estimated by $\boldsymbol{K_d} = \frac{\textbf{number of frames in unbound state}}{\textbf{number of frames in bound state}}$ for each histone type. Then, the histone tail's binding free energy with DNA ($\Delta G_0$ in kcal/mol) is derived from $\mathbf{K_d}$ using the following equation: $\boldsymbol{\Delta G_0 = RT \ln K_d}$. For each simulation run, the number of observed unbinding events of each tail copy was counted and only included runs that had at least five unbinding events for the analysis.

| Residue | Node degree (WT) | Node degree (G33R) | Node degree (G34R) |
|---|---|---|---|
| A1 | 0.126 | 0.317 | 0.035 |
| R2 | 0.291 | 0.525 | 0.382 |
| T3 | 0.337 | 0.465 | 0.214 |
| K4 | 0.21 | 0.267 | 0.188 |
| Q5 | 0.464 | 0.663 | 0.609 |
| T6 | 0.377 | 0.428 | 0.409 |
| A7 | 0.333 | 0.269 | 0.27 |
| R8 | 0.691 | 0.482 | 0.573 |
| K9 | 0.291 | 0.414 | 0.394 |
| S10 | 0.639 | 0.491 | 0.424 |
| T11 | 0.467 | 0.639 | 0.432 |
| G12 | 0.268 | 0.247 | 0.233 |
| G13 | 0.318 | 0.311 | 0.273 |
| K14 | 0.38 | 0.41 | 0.405 |
| A15 | 0.503 | 0.533 | 0.38 |
| P16 | 0.351 | 0.32 | 0.274 |
| R17 | 0.84 | 0.953 | 0.57 |
| K18 | 0.387 | 0.466 | 0.494 |
| Q19 | 0.67 | 0.923 | 0.861 |
| L20 | 0.671 | 1.081 | 0.699 |
| A21 | 0.554 | 0.832 | 0.49 |
| T22 | 0.626 | 0.919 | 0.747 |
| K23 | 0.456 | 0.736 | 0.465 |
| A24 | 0.296 | 0.66 | 0.409 |
| A25 | 0.316 | 0.534 | 0.423 |
| R26 | 0.699 | 0.791 | 0.553 |
| K27 | 0.384 | 0.538 | 0.243 |
| S28 | 0.43 | 0.846 | 0.312 |
| A29 | 0.315 | 0.705 | 0.384 |
| P30 | 0.598 | 0.763 | 0.398 |
| S31 | 0.641 | 0.949 | 0.531 |
| T32 | 0.539 | 0.838 | 0.442 |
| G33 | 0.344 | 1.115 | 0.339 |
| G34 | 0.458 | 0.554 | 0.575 |
| V35 | 0.384 | 0.286 | 0.295 |
| K36 | 0.435 | 0.493 | 0.338 |
| K37 | 0.141 | 0.115 | 0.176 |

**Supplementary Table 3.** The degree of nodes for each tail residue from the interaction graph generated from the structural ensembles in simulations using the RING 4.0 sever.

| System | WT | G33R mutant | G34R mutant |
|---|---|---|---|
| VDW | THR32-ALA29: 0.0741 | ARG33-PRO30: 0.1515 | ARG34-SER31: 0.0851 |
| | GLY34-SER31: 0.0731 | THR32-ALA29: 0.1231 | GLY33-PRO30: 0.0547 |
| | THR32-VAL35: 0.0492 | ARG33-LYS36: 0.0876 | THR32-ALA29: 0.0438 |
| | GLY33-PRO30: 0.0425 | ARG33-ALA29: 0.0855 | ARG34-PRO30: 0.0304 |
| | GLY33-LYS36: 0.0197 | GLY34-SER31: 0.0788 | VAL35-THR32: 0.0258 |
| | THR32-SER28: 0.0181 | THR32-SER28: 0.0690 | ARG34-GLN5: 0.0237 |
| H-Bond | GLY34(D)-SER31(A): 0.1472 | THR32(D)-SER28(A): 0.1947 | ARG34(D)-SER31(A): 0.1088 |
| | THR32(D)-ALA29(A): 0.1047 | THR32(D)-ALA29(A): 0.1700 | GLY33(D)-PRO30(A): 0.0825 |
| | GLY33(D)-PRO30(A): 0.0684 | ARG33(D)-ALA29(A): 0.1669 | THR32(D)-ALA29(A): 0.0794 |
| | GLY33(D)-ALA29(A): 0.0383 | ARG33(D)-PRO30(A): 0.1643 | THR32(A)-VAL35(D): 0.0304 |
| | THR32(A)-VAL35(D): 0.0358 | GLY34(D)-PRO30(A): 0.1489 | ARG34(D)-GLN5(A): 0.0294 |
| | GLY33(A)-LYS36(D): 0.0244 | GLY34(D)-SER31(A): 0.1087 | THR32(D)-SER28(A): 0.0242 |

**Supplementary Table 4.** Hydrogen bonds (H-bond) and van der Waals (VDW) interactions observed between the hinge region (TGG) and other H3.3 tail residues. The probability score for each interaction represents the fraction of MD trajectories in which the interaction was observed. In the H-bond analysis, “D” denotes the donor residue and “A” denotes the acceptor residue.

**a**

```
H3.3    artkqtarkstggkaprkqlatkaarksapstggvkkphryrpgtvalreirryqkstel  60
H3.1    artkqtarkstggkaprkqlatkaarksapatggvkkphryrpgtvalreirryqkstel  60
H3.2    artkqtarkstggkaprkqlatkaarksapatggvkkphryrpgtvalreirryqkstel  60
        ******************************.*****************************

H3.3    lirklpfqrlvreiaqdfktdlrfqsaaigalqeaseaylvglfedtnlcaihakrvti 119
H3.1    lirklpfqrlvreiaqdfktdlrfqssavmalqeaceaylvglfedtnlcaihakrvti 119
H3.2    lirklpfqrlvreiaqdfktdlrfqssavmalqeaseaylvglfedtnlcaihakrvti 119
        **************************:*: *****.***********************

H3.3    mpkdiqlarrirgera     135
H3.1    mpkdiqlarrirgera     135
H3.2    mpkdiqlarrirgera     135
        ****************
```

**b**

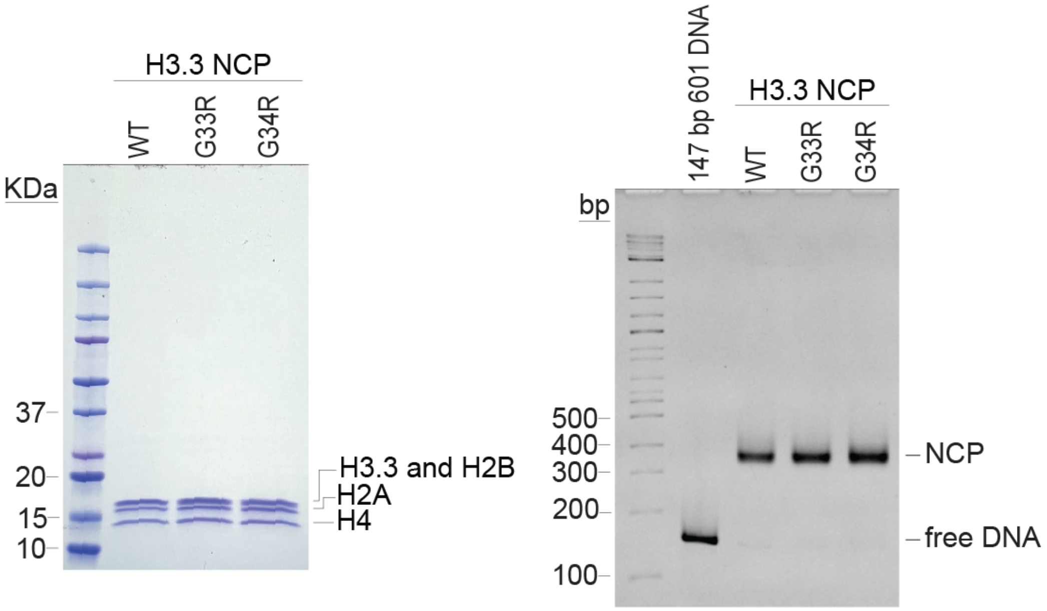


**Supplementary Figure 1. (a)** Alignment of histone H3 variant sequences H3.1 (NP_003522.1), H3.2 (NP_066403.2), H3.3 (P84243.2) aligned using Clustal Omega. **(b)** SDS-PAGE (left) and Native-PAGE (right) analysis of the wild-type (WT) and G33R, G34R mutants. The 147bp 601 sequence is shown as reference in the native-PAGE.

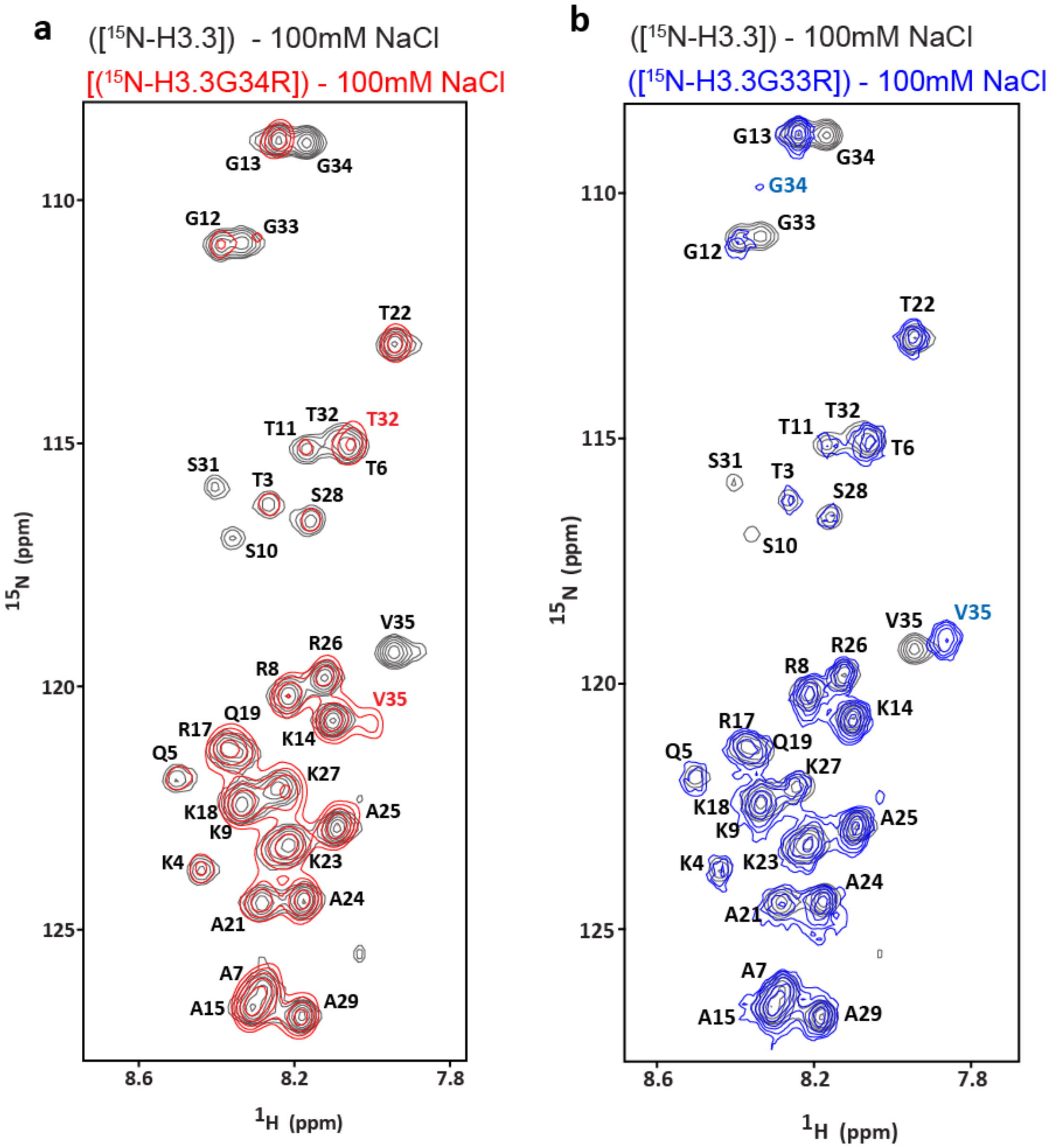


**Supplementary Figure 2. (a)** Overlay of the $^1$H, 15N-HSQC spectra for wild-type nucleosomes containing $^{15}$N-enriched H3.3 ([$^{15}$N-H3.3]) shown in black or the H3.3G34R mutant ([$^{15}$N-H3.3G34R]) shown in red. **(b)** Overlay of the $^1$H, 15N-HSQC spectra for wild-type nucleosomes containing $^{15}$N-enriched H3.3 ([$^{15}$N-H3.3]) shown in black or the H3.3G33R mutant ([$^{15}$N-H3.3G33R]) shown in blue. All spectra were collected in the presence of 100mM NaCl.

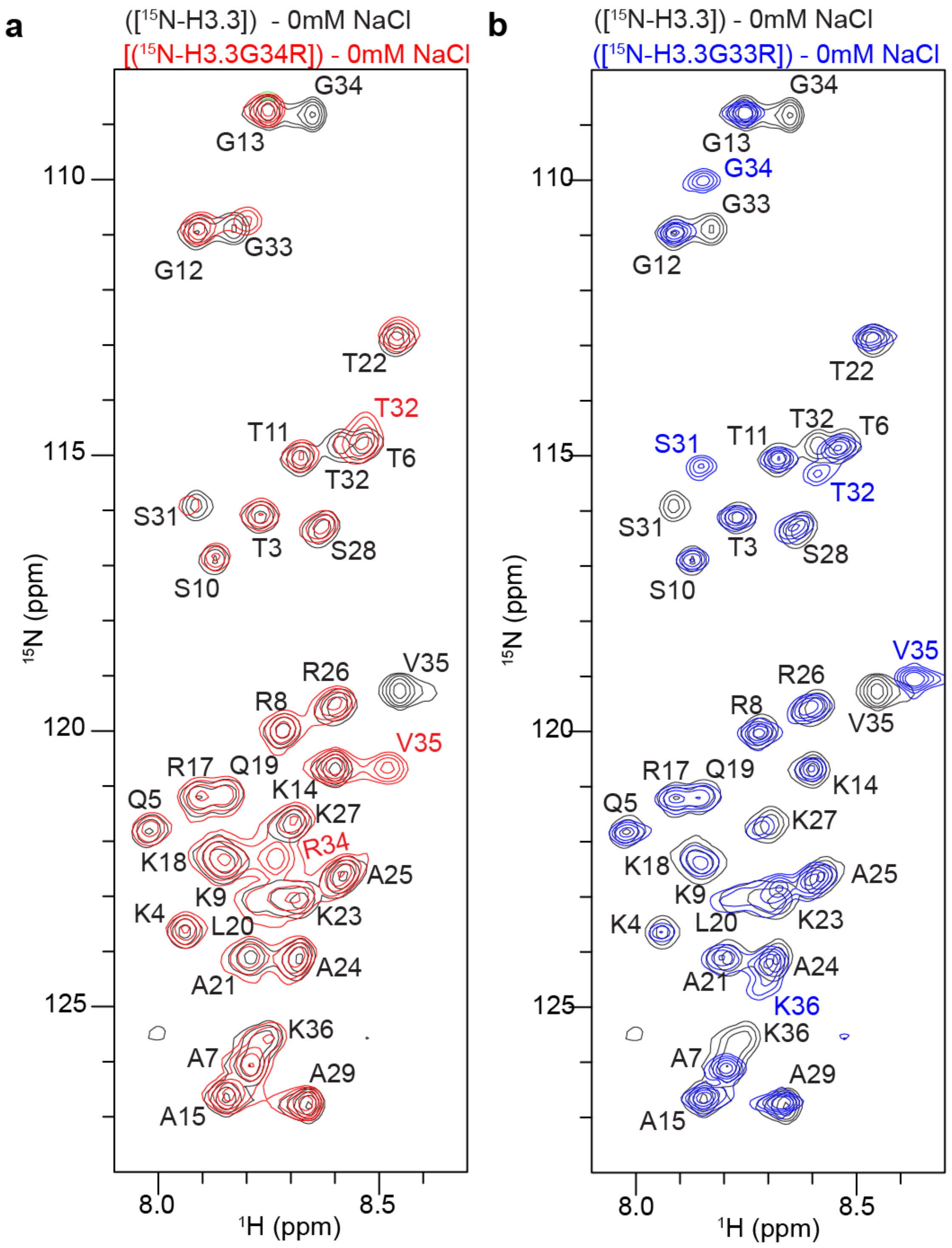


**Supplementary Figure 3. (a)** Overlay of the $^{1}H$, 15N-HSQC spectra for wild-type nucleosomes containing $^{15}N$-enriched H3.3 ([$^{15}N$-H3.3]) shown in black or the H3.3G34R mutant ([$^{15}N$-H3.3G34R]) shown in red. **(b)** Overlay of the $^{1}H$, 15N-HSQC spectra for wild-type nucleosomes containing $^{15}N$-enriched H3.3 ([$^{15}N$-H3.3]) shown in black or the H3.3G33R mutant ([$^{15}N$-H3.3G33R]) shown in blue. All spectra were collected in the presence of 0mM NaCl.

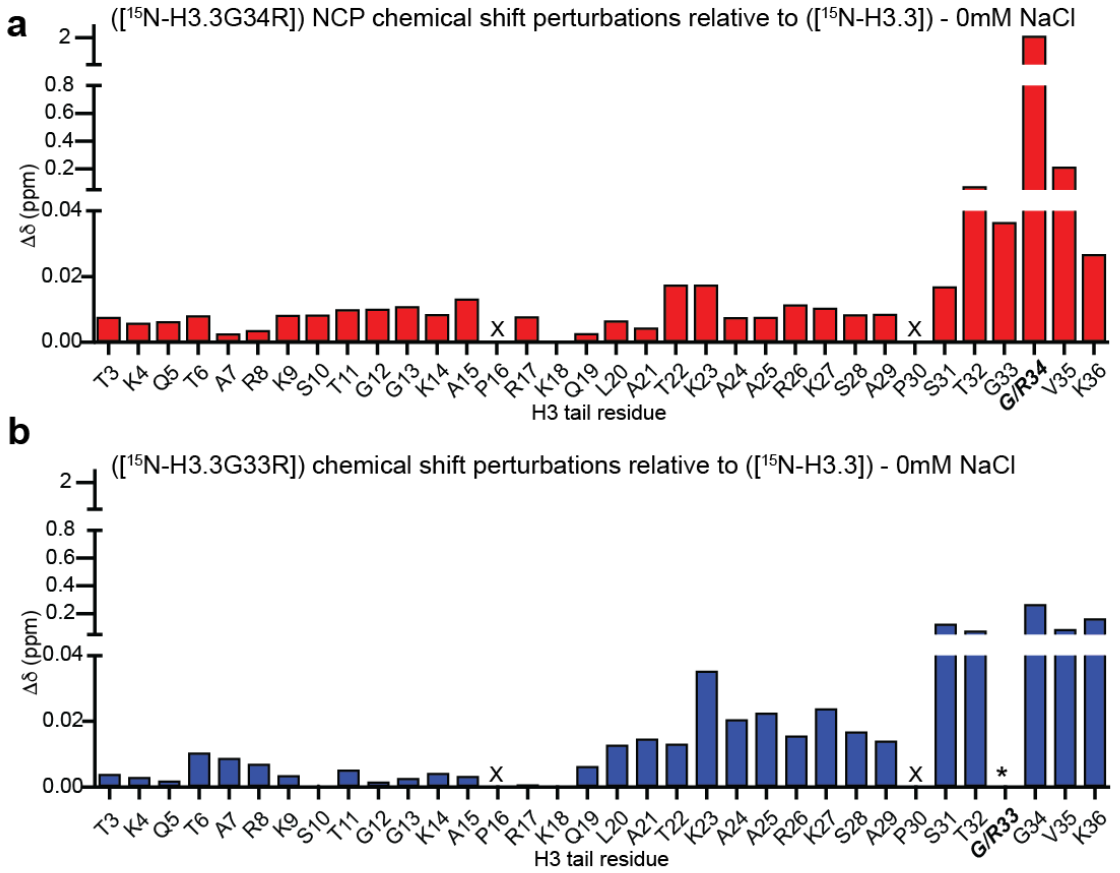


**Supplementary Figure 4.** Normalized chemical shift changes (Δδ) between $^{1}$H,$^{15}$N-HSQC spectra plotted as a function of H3.3 tail residues between WT H3.3 and H3.3G34R containing nucleosomes (**a**, red) or WT H3.3 and H3.3G33R containing nucleosomes (**b**, blue). X denotes a proline residue which does not have a backbone amide and thus no corresponding peak, * denotes a missing peak. All data were collected at 0mM NaCl.

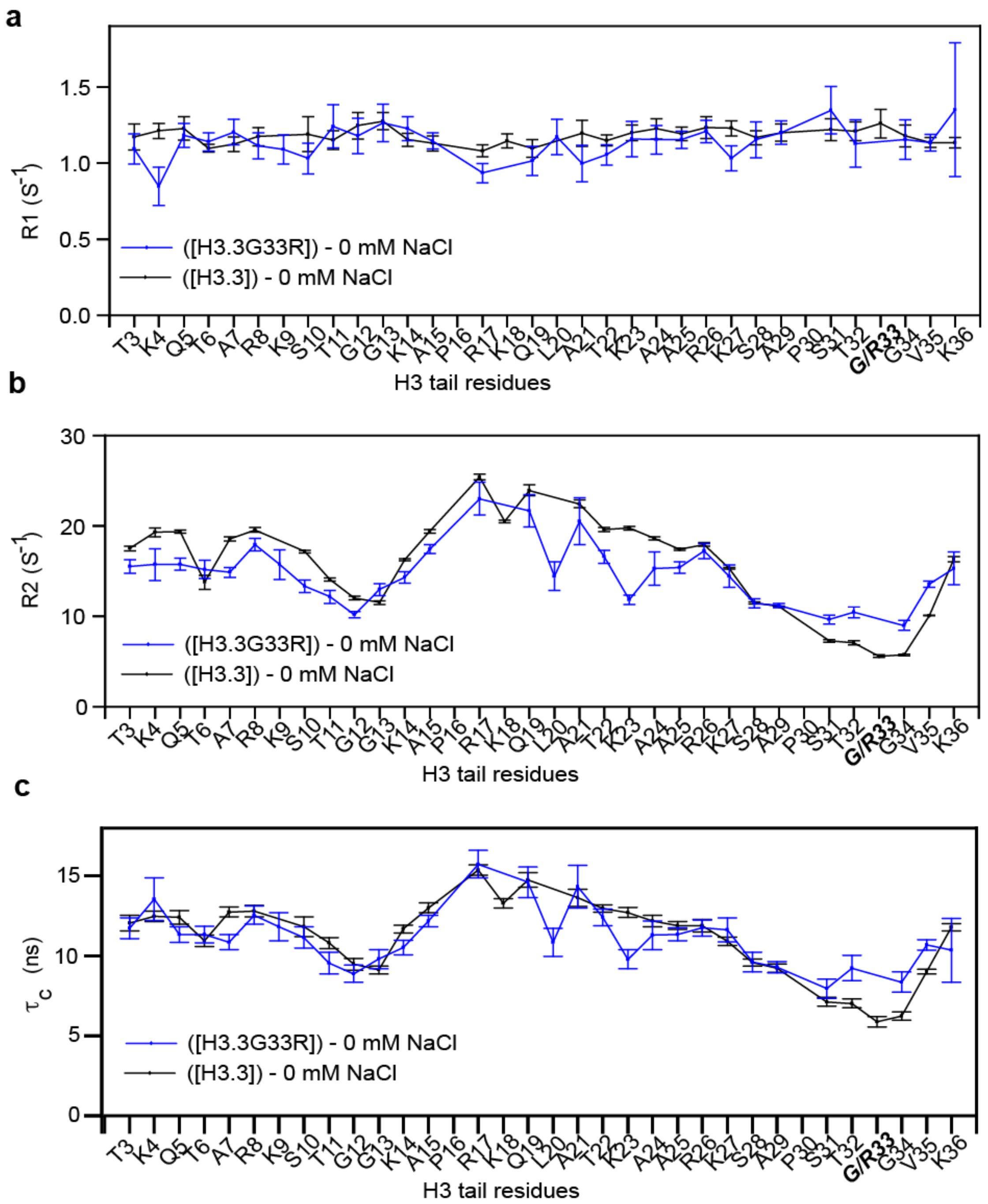


**Supplementary Figure 5.** $^{15}$N-relaxation rates R1 **(a)**, R2 **(b)**, and computed correlation times (**c**, $\tau_C$) as a function of H3.3 tail residue for nucleosomes containing WT H3.3 (black) or mutant H3.3G33R (blue) All data was collected at 0mM NaCl.

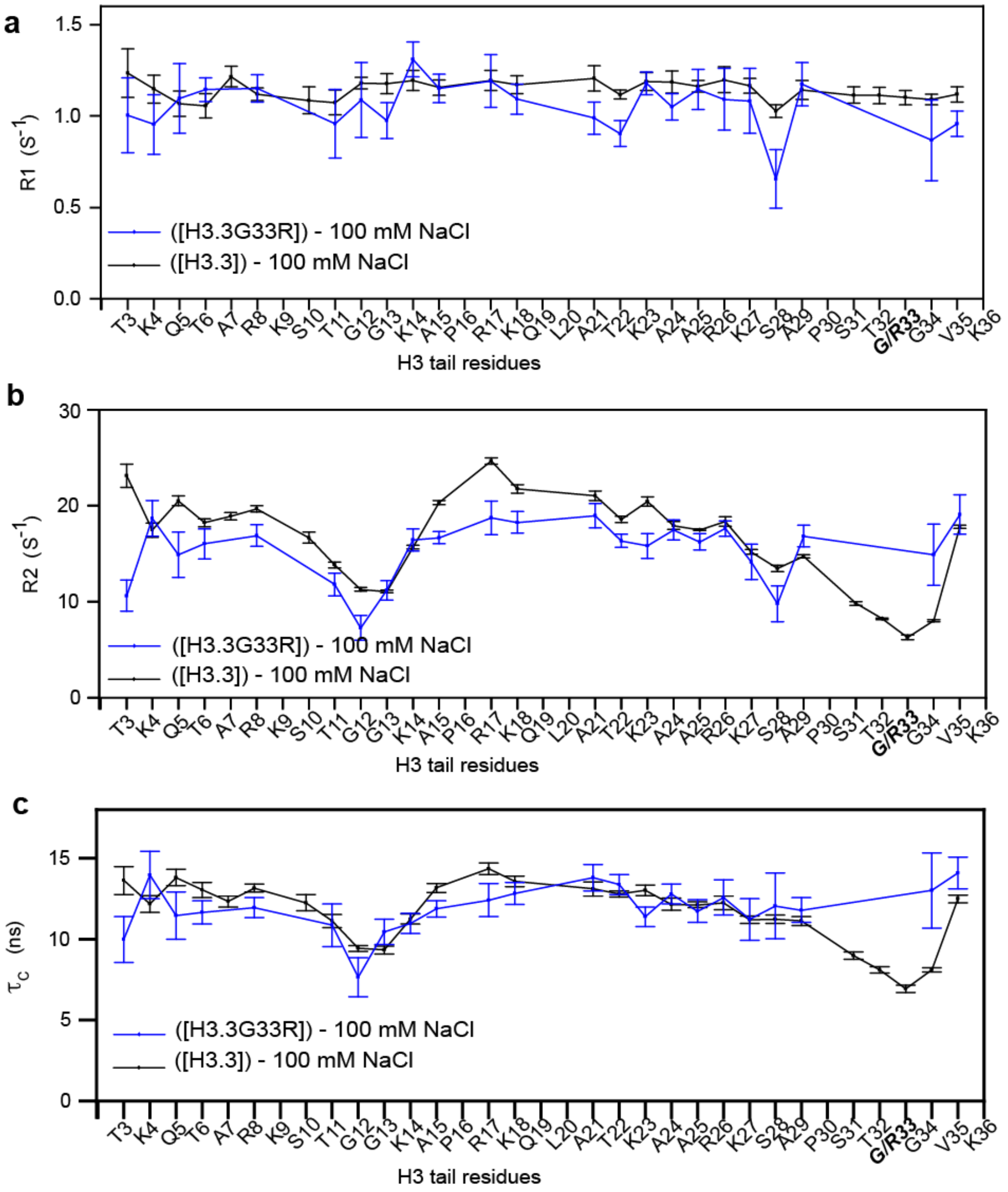


**Supplementary Figure 6.** $^{15}$N-relaxation rates R1 **(a)**, R2 **(b)**, and computed correlation times (**c**, $\tau_C$) as a function of H3.3 tail residue for nucleosomes containing WT H3.3 (black) or mutant H3.3G33R (blue) All data was collected at 100mM NaCl.

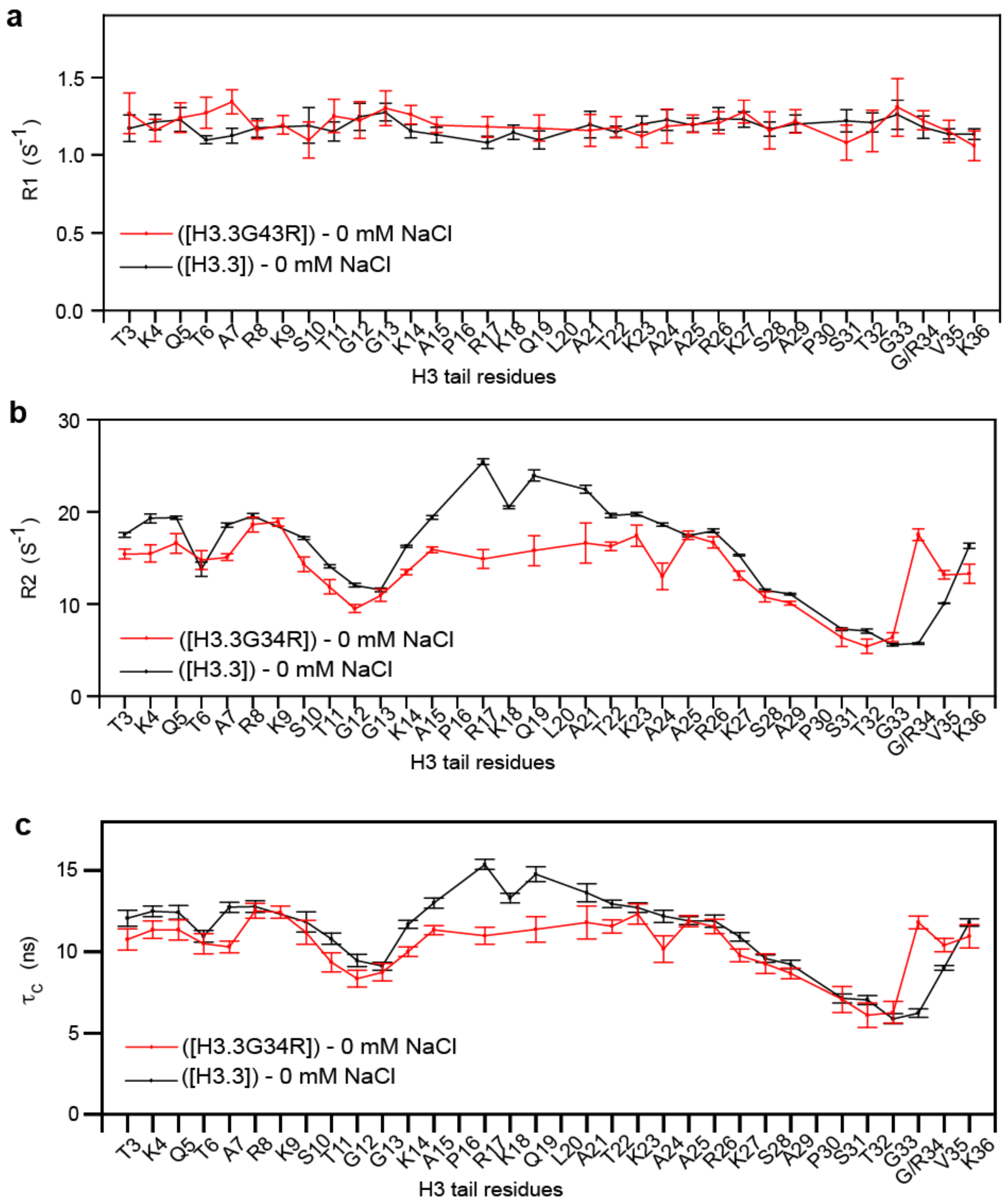


**Supplementary Figure 7.** $^{15}$N-relaxation rates R1 **(a)**, R2 **(b)**, and computed correlation times (**c**, $\tau_C$) as a function of H3.3 tail residue for nucleosomes containing WT H3.3 (black) or mutant H3.3G34R (red). All data was collected at 0mM NaCl.

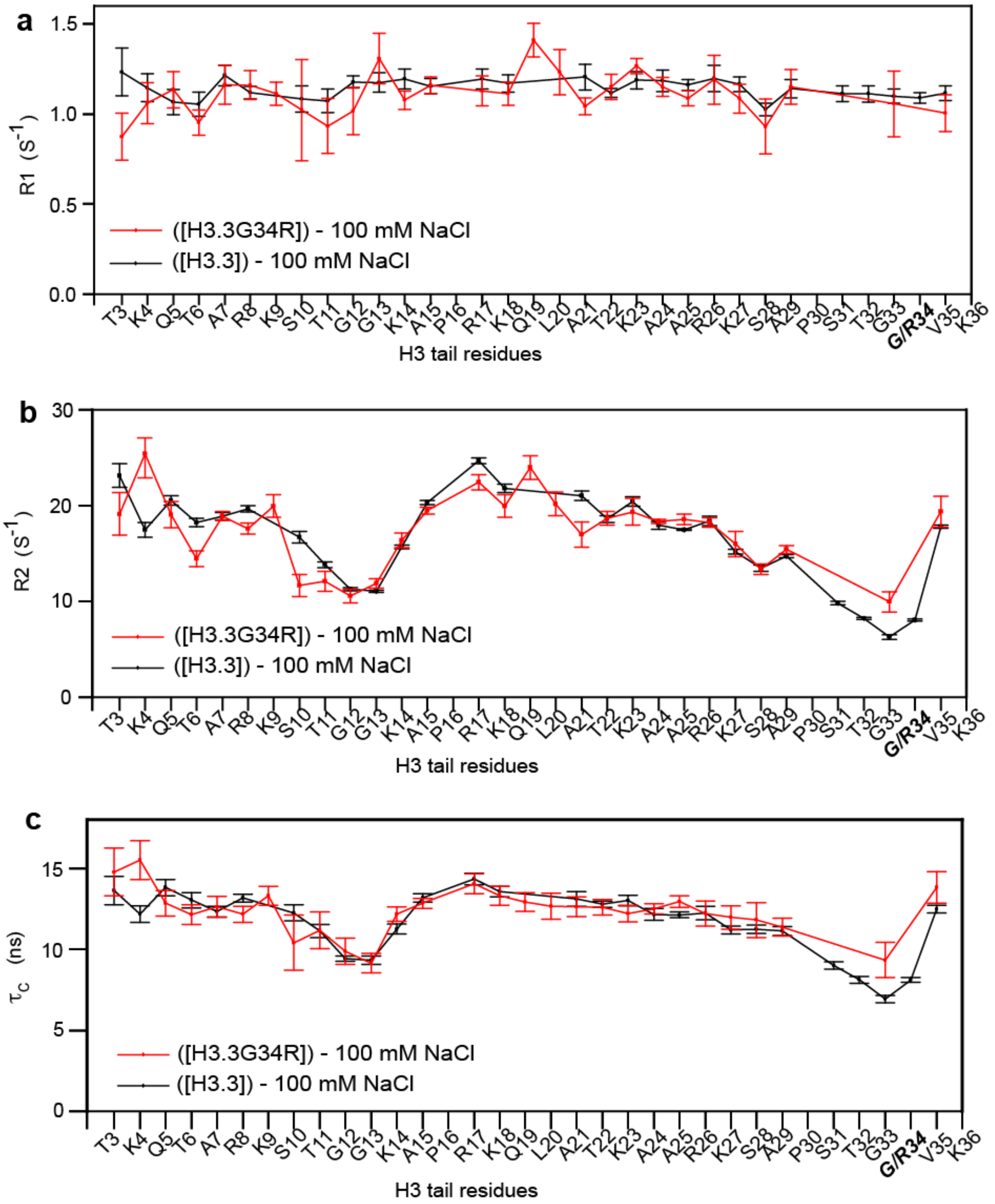


**Supplementary Figure 8.** $^{15}$N-relaxation rates R1 **(a)**, R2 **(b)**, and computed correlation times (**c**, $\tau_C$) as a function of H3.3 tail residue for nucleosomes containing WT H3.3 (black) or mutant H3.3G34R (red). All data was collected at 100mM NaCl.

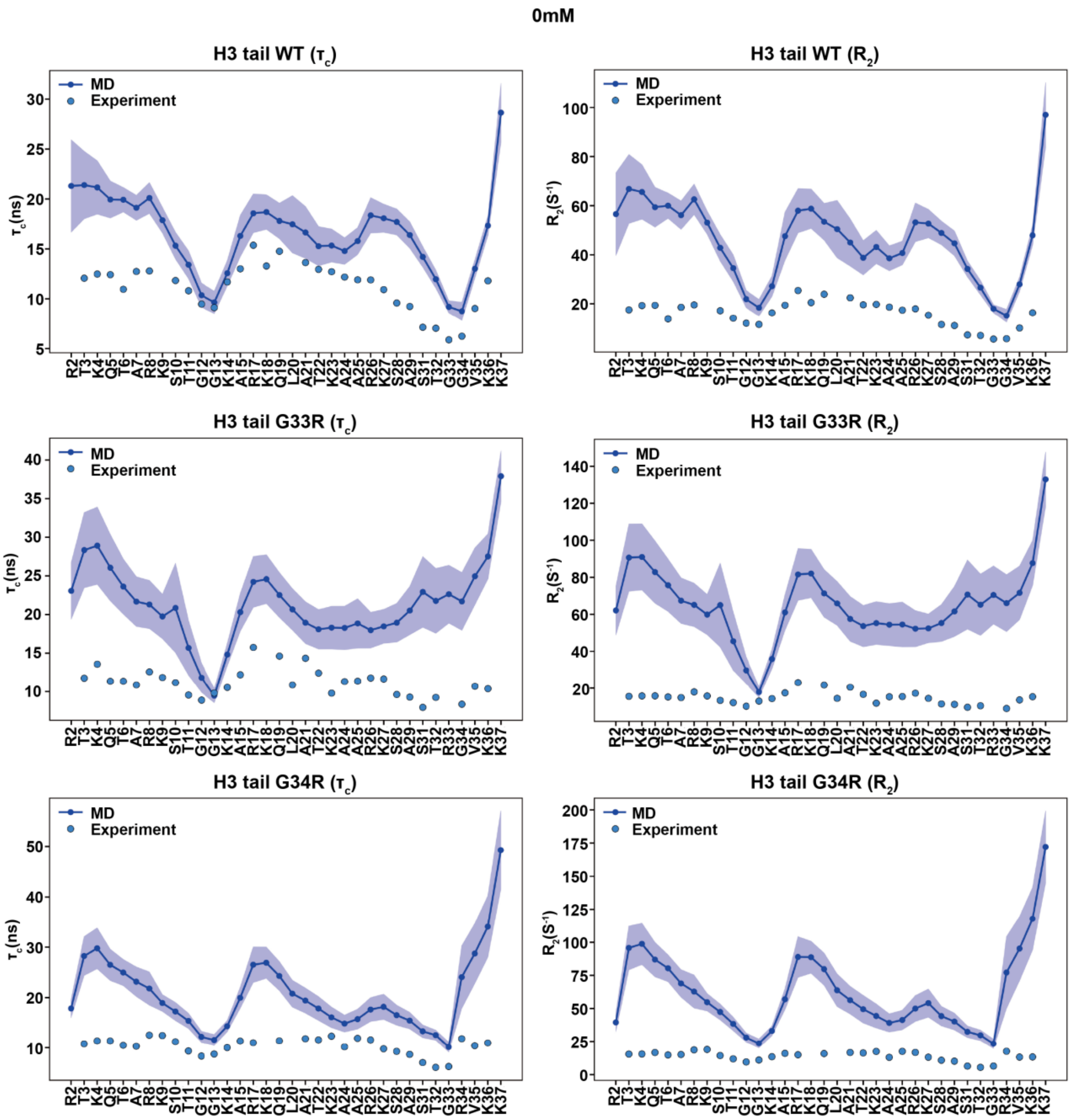


**Supplementary Figure 9.** Comparison of predicted $\tau_c$ and $R_2$ values with experimental data measured at 0mM NaCl of wild-type, G33R and G34R H3 tails.

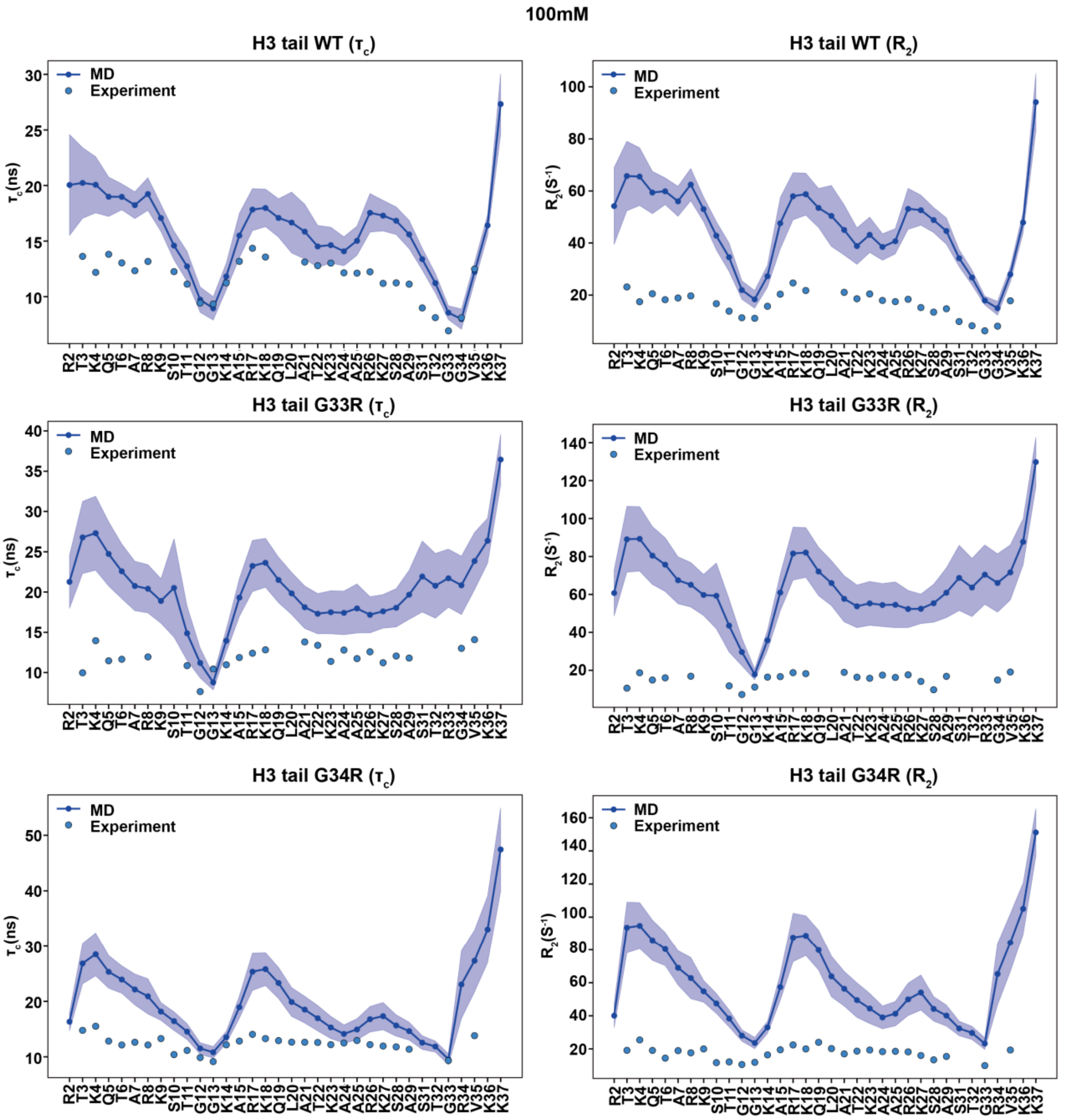


**Supplementary Figure 10.** Comparison of predicted $\tau_c$ and $R_2$ values with experimental data measured at 100mM NaCl of wild-type, G33R and G34R H3 tails.

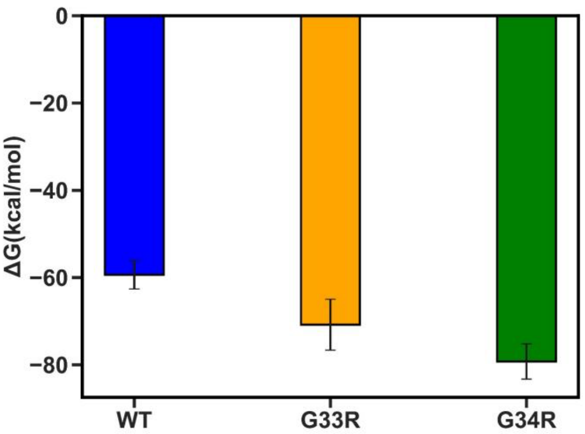


**Supplementary Figure 11.** H3.3 tail-DNA binding free energies calculated using the MM/GBSA approach. The reported values are averaged over 1 ns frames and the error bars represent the standard errors of the mean calculated from 10 independent measurements.

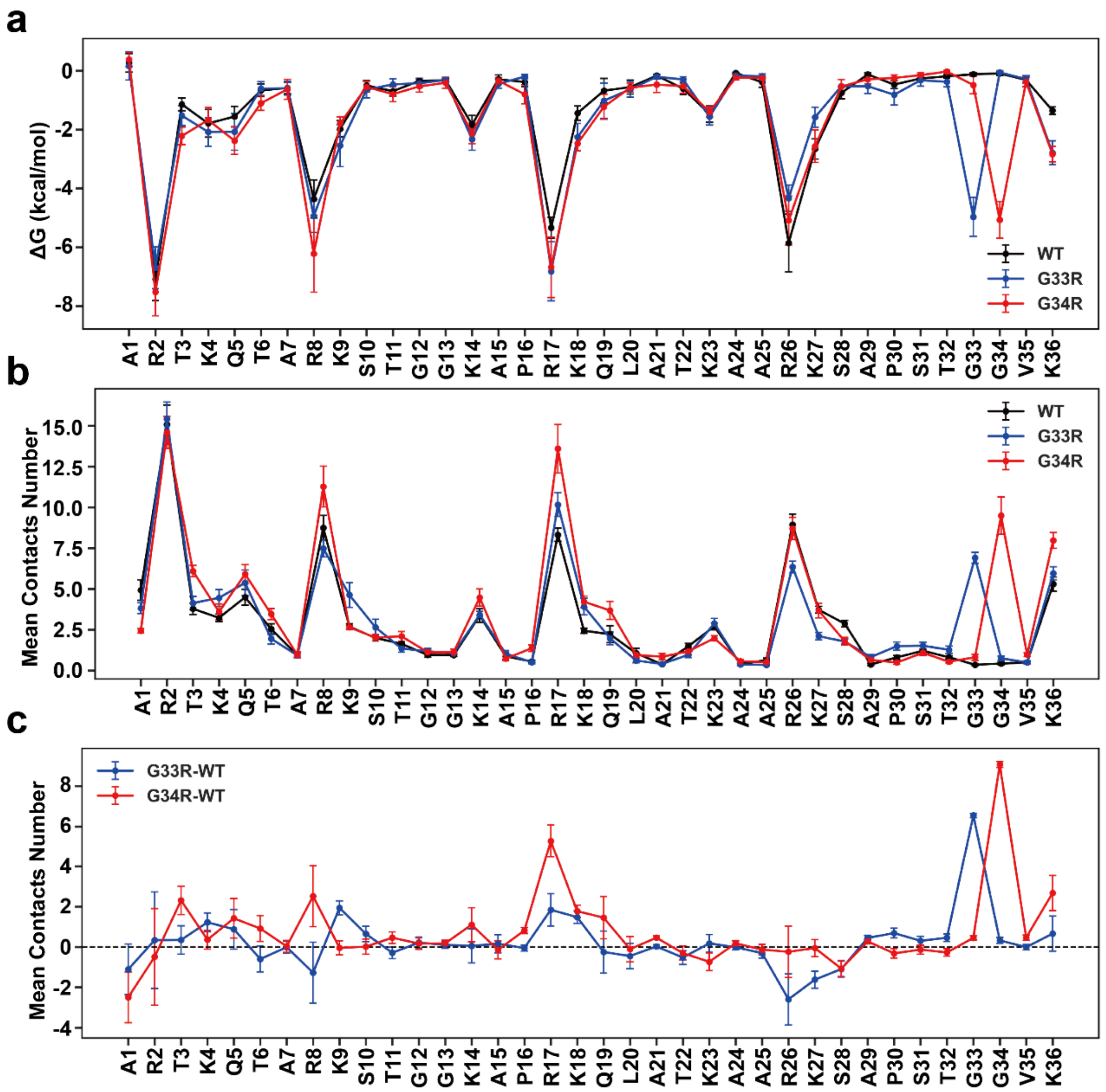


**Supplementary Figure 12. (a)** Binding free energies of wild-type and mutant H3.3 tails plotted against tail residue numbers calculated from MM/GBSA approach. Binding free energies are averaged over 1 ns frames, and the error bars represent standard errors of the mean calculated from independent simulation runs. **(b)** Mean number of contacts between tail residues and DNA molecules in WT and mutated H3.3 tails. **(c)** The change of the mean number of contacts between tail residues and DNA molecules upon H3.3G33R and H3.3G34R mutants.

a

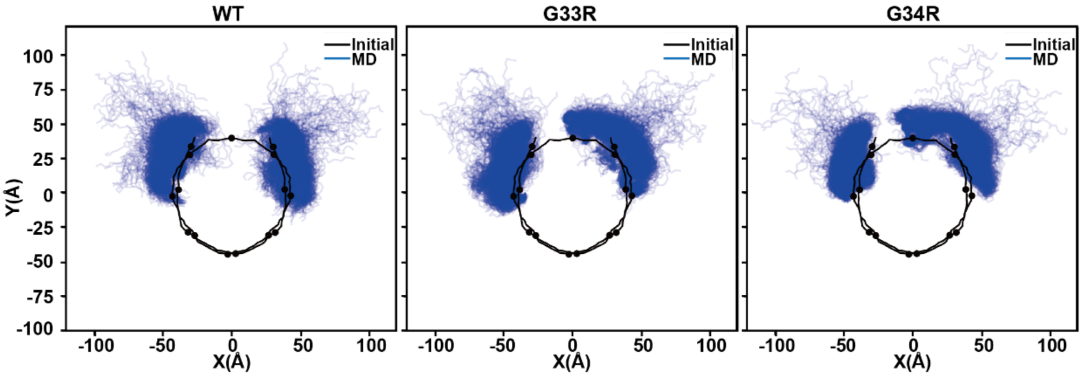


**Supplementary Figure 13.** Projection of H3.3 tail conformations in simulation runs of WT H3.3, H3.3G33R, and H3.3G34R nucleosomes. The H3 tail conformational ensembles were projected onto the plane perpendicular to the superhelical axis for MD snapshots. The black line and dots indicate the integer and half-integer SHL values of the initial conformation of DNA molecules.

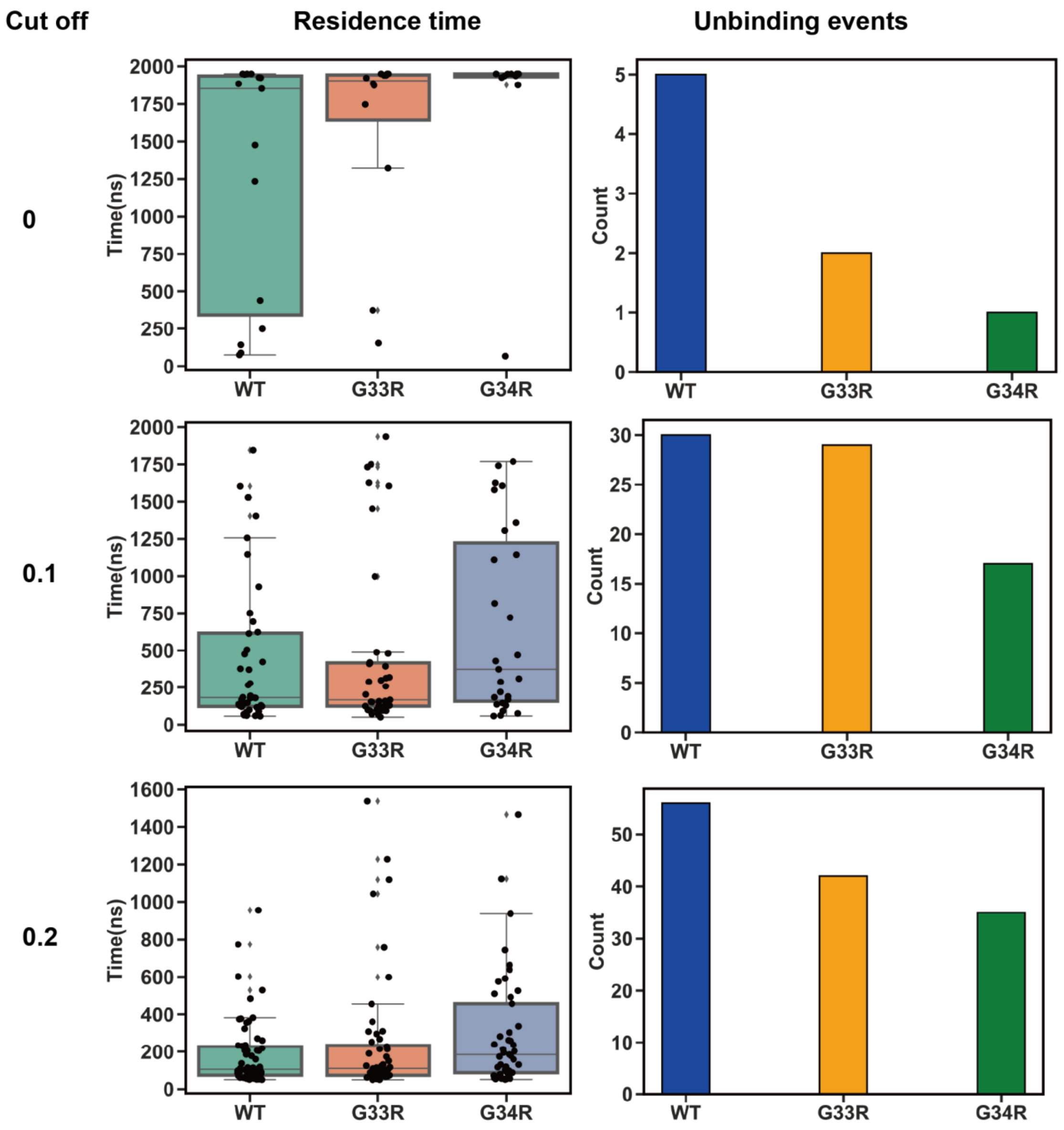


**Supplementary Figure 14.** Dependence of histone H3.3 tail residence time ($\tau_f$) on the chosen cut-off values for the percentage of frames which maintained contacts with DNA in simulations (noted on the left) for the full tail unbound state definition. An unbound state is defined if the percentage of tail residues maintaining contacts with DNA is less than the chosen cut-off. Box-plot elements are defined as: center line, median; box limits, upper and lower quartiles; whiskers are drawn at values equal to 1.5× interquartile range. Unbinding events were summed across all simulation runs, and events with residence times shorter than 50 ns were excluded from the analysis.

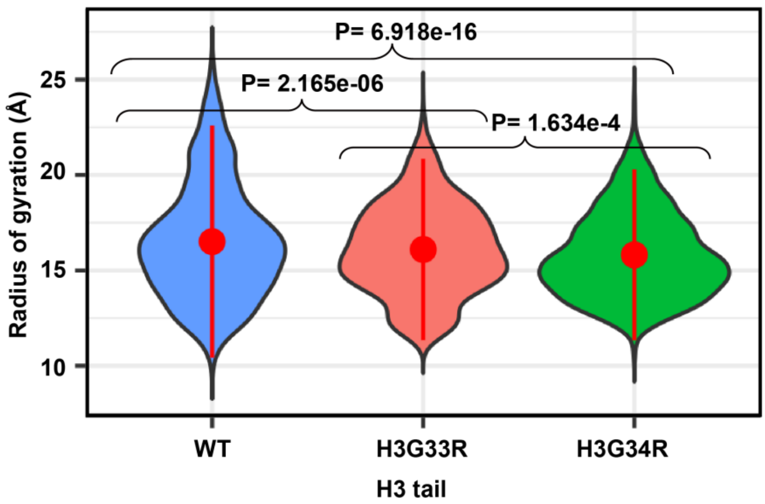


**Supplementary Figure 15.** Comparison of the calculated radius of gyration (RG) of the H3.3 tail between WT and mutated tails.

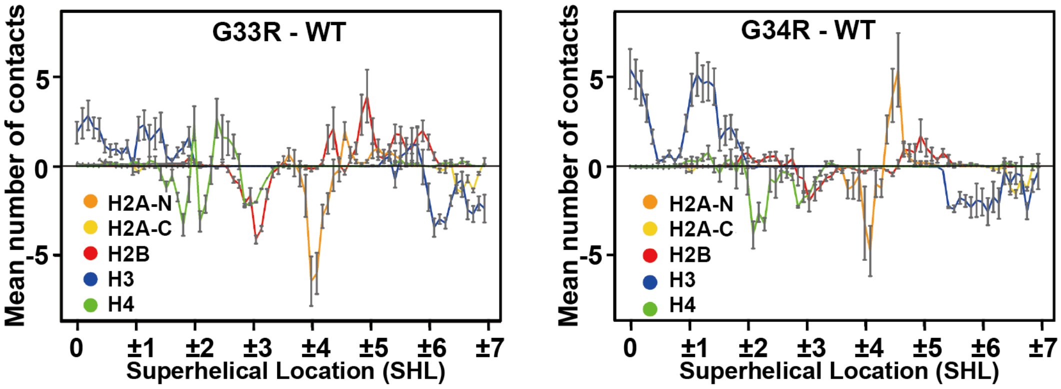


**Supplementary Figure 16**. The change in the mean number of contacts between each histone tail and nucleosomal DNA upon H3.3G33R or H3.3G34R mutations. For each mutation, the change of the mean number of contacts is calculated as the mean number of contacts between DNA and mutant tails minus the mean number of contacts between DNA and WT tails. The reported values are averaged, and the error bars represent the standard errors of the mean for mutant tails calculated from independent simulation runs (n=10).

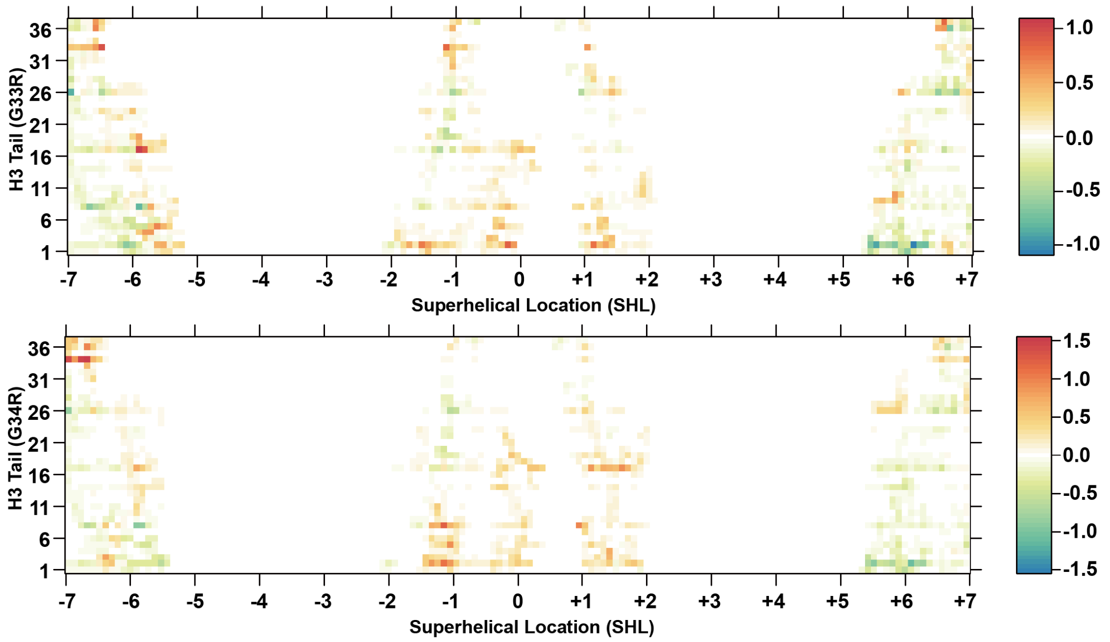


**Supplementary Figure 17.** A heat plot of the change in the mean number of tail-DNA contacts for mutant H3.3G33R or H3.3G34R tails plotted against thesuper-helical location (SHL). For each mutation, the change in the mean number of contacts is calculated as the mean number of contacts between DNA base pairs and mutant tail residues minus the mean number of contacts between DNA base pairs and WT tail residues. Red indicates an increase in the mean numbers of contacts.
